# Assessment of The Broad-Spectrum Host Targeting Antiviral Efficacy of Halofuginone Hydrobromide in Human Airway, Intestinal and Brain Organoid Models

**DOI:** 10.1101/2023.11.01.565121

**Authors:** Inés García-Rodríguez, Giulia Moreni, Pamela E. Capendale, Lance Mulder, Ikrame Aknouch, Renata Vieira de Sá, Nina Johanneson, Eline Freeze, Hetty van Eijk, Gerrit Koen, Katja Wolthers, Dasja Pajkrt, Adithya Sridhar, Carlemi Calitz

## Abstract

Halofuginone hydrobromide has shown potent antiviral efficacy against a variety of viruses such as SARS-CoV-2, dengue, or chikungunya virus, and has, therefore, been hypothesized to have broad-spectrum antiviral activity. In this paper, we tested this broad-spectrum antiviral activity of Halofuginone hydrobomide against viruses from different families (*Picornaviridae, Herpesviridae, Orthomyxoviridae, Coronaviridae,* and *Flaviviridae).* To this end, we used relevant human models of the airway and intestinal epithelium and regionalised neural organoids. Halofuginone hydrobomide showed antiviral activity against SARS-CoV-2 in the airway epithelium with no toxicity at equivalent concentrations used in human clinical trials but not against any of the other tested viruses.

**Graphical abstract:** 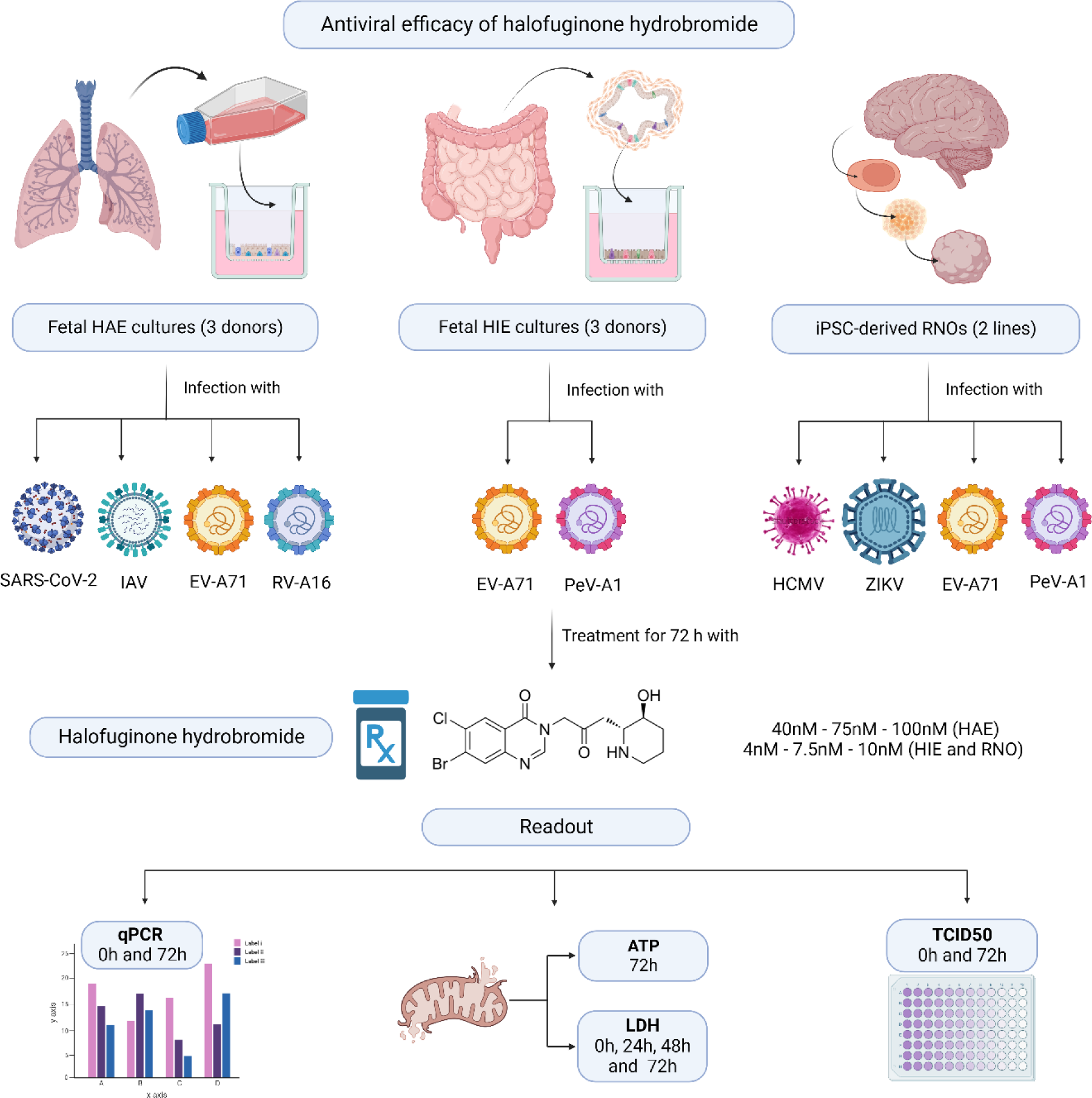

**Highlights:**

- Halofuginone hydrobromide was identified as a possible broad-spectrum host targeting antiviral drug.
- Human organoid models offer a physiologically relevant and clinically translatable model for antiviral research.
- Halofuginone hydrobromide shows antiviral efficacy against SARS-CoV-2, but not against EV-A71, PeV-A1, IAV, RV-A16, HCMV or ZIKV in relevant organoid models.
- The efficacy of Halofuginone hydrobromide is concentration dependent as well as on proline content of the host receptor(s) or host factors for the specific virus in question.

## 1 Introduction

The devastating consequences that COVID-19 had on mortality, morbidity, and the social economic sphere has proven that a novel approach in how we deal with future events is essential (British-Academy, 2021). One possible solution is the identification and utilization of broad-spectrum antivirals. However, effective prophylactic and antiviral treatment options with broad-spectrum action remain elusive. COVID-19 was a driving force in shifting towards the repurposing of FDA approved drugs as potentially new antiviral compounds. One such compound, halofuginone hydrobromide, was identified as an antiviral targeting host factors with predicted activity against multiple viruses (Chitalia and Munawar, 2020; Geraghty et al., 2021; Hwang et al., 2019).

Halofuginone hydrobromide (HH) {7-bromo-6-chloro-3-[3-(3-hydroxy-2-piperidinyl)-2- oxopropyl] -4(3H)-quinazolinone}, a febrifugine analog, is an alkaloid isolated from the roots of the *Dichroa febrifuga* Lour., (Hydrangeaceae) plant, commonly known as Chinese quinine. Originally HH was used as an additive to poultry feed, however, in recent years it has shown promise as an effective antifibrotic, anti-inflammatory, antimalarial and anti-cancer drug (Keller et al., 2012; Pines and Spector, 2015). The proposed broad-spectrum antiviral activity of HH is attributed to the molecules ability to competitively bind to and inhibit prolyl-tRNA synthetase (Hwang et al., 2019). Upon HH binding to the prolyl tRNA synthetase there is an accumulation of uncharged prolyl tRNAs. This results in the suppression of bulk translation through amino acid starvation response (AAR) (Keller et al., 2012). In a 2019 study by Hwang and colleagues, HH showed efficacy against both chikungunya virus (CHIKV) and dengue virus (DENV) (Hwang et al., 2019). While in 2021, Chen and colleagues showed the efficacy of HH against SARS-CoV-2 viral entry and replication in both cell lines and human bronchial airway cultures (Chen et al., 2021). The results suggest that HH could be a possible broad-spectrum antiviral drug. This could either be directly against the virus by direct inhibition of viral translation or indirectly by affecting the host translation, for example, by preventing the correct expression of cellular receptors necessary for the virus to enter the cell.

With the current study, we evaluated the proposed broad spectrum antiviral efficacy of HH in human organotypic models for a range of viruses. Human organotypic models have emerged as valuable tools in studying viral infections and antiviral drug screening (Masmoudi et al., 2023). These cultures closely mimic the cell composition and function of their respective *in vivo* tissues, providing a physiologically relevant platform to investigate host-pathogen interactions and therapeutic interventions (Krenn et al., 2021; Saxena et al., 2016). As a result of these advantages, human organotypic cultures are expected to perform better than 2D cell line models and animal models which have 95% drug attrition rates (Richman and Nathanson, 2016) and translational challenges (Sridhar et al., 2020).

To this end, we assessed the toxicity and antiviral potential of HH against a panel of viruses: enterovirus A71 (EV-A71), parechovirus A1 (PeV-A1), rhinovirus A16 (RV-A16), influenza A virus (IAV), human cytomegalovirus (HCMV), Zika virus (ZIKV), and severe acute respiratory syndrome coronavirus 2 (SARS-CoV-2), utilizing human intestinal, airway, and brain models. The panel of viruses was chosen due to their significant impact on human health, their diverse tropism, and the pressing need for new therapeutic approaches (Supplementary Table 1). The choice of models used for HH efficacy assessment was based on the human primary and secondary replication sites for each of the aforementioned viruses (Supplementary Table 1 and Figure 1). This comprehensive evaluation will provide insights into the potential of HH as a host targeting broad-spectrum antiviral agent.

**Figure 1.**
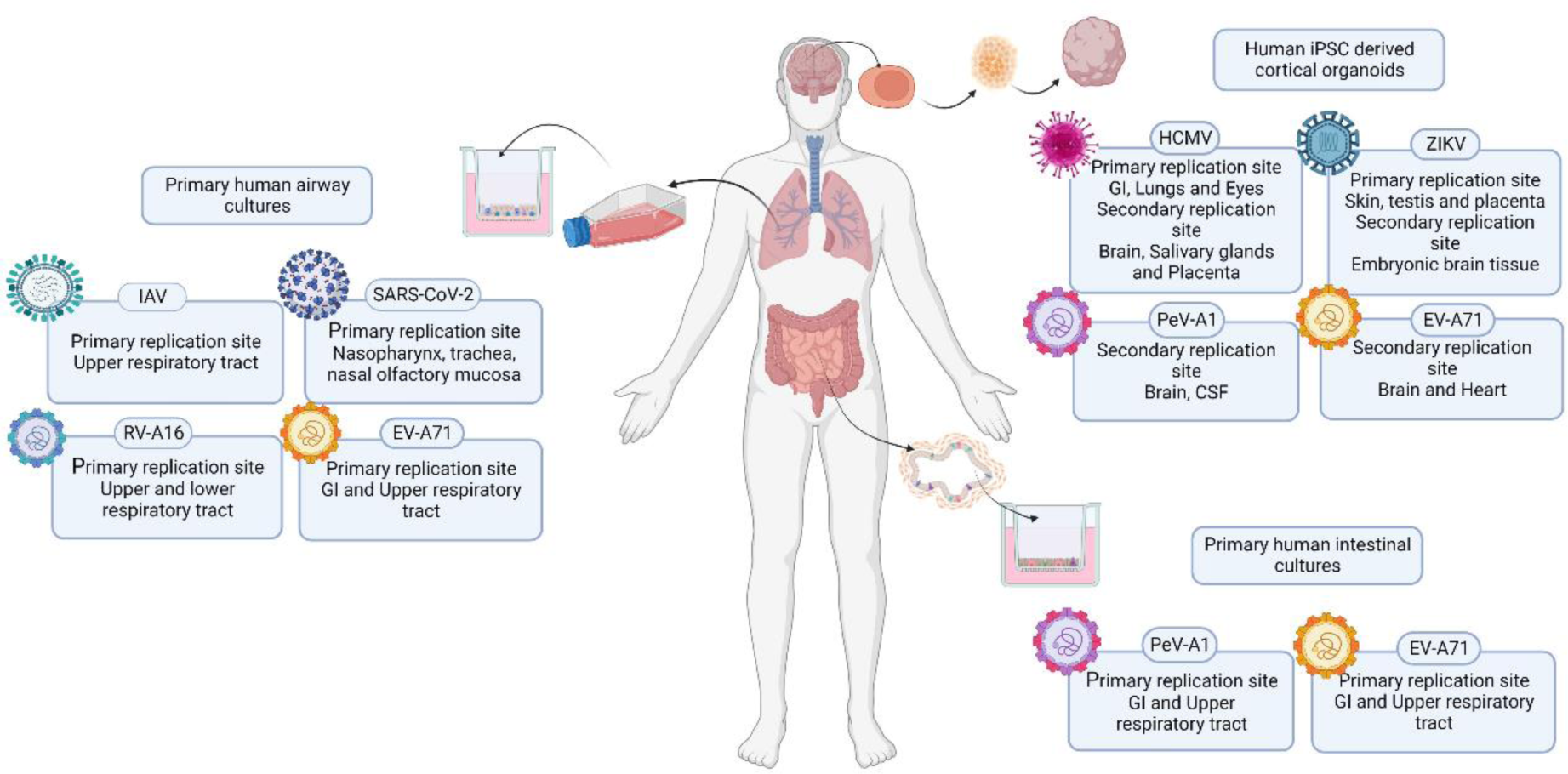
Schematic representation of the study, with the studied viruses and organotypic models used. GI: gastrointestinal tract, CSF: cerebrospinal fluid, iPSC: induced pluripotent stem cells.

## 2 Materials and methods

### 2.1 Ethics

Human fetal intestinal and airway tissue, gestational age 16–20 weeks, was obtained from a private clinic by the HIS Mouse Facility of the Amsterdam University Medical Centers. All donors supplied written informed consents for the use of fetal material anonymized for research purposes. The use of the anonymized material for medical research purposes is covered by a Dutch law (Wet foetaal weefsel and Article 467 of Behandelingsovereenkomst).

### 2.2 Cell lines and virus strains

Human colorectal adenocarcinoma (HT-29) cells (HTB-38^™^ ATCC), human rhabdomyosarcoma (RD) cells (CCL-136^™^ ATCC), African green monkey (*Chlorocebus aethiops*) kidney (Vero) cells (kindly provided by the National Institute of Public Health and the Environment (RIVM) the Netherlands), Vero E6 cells (CRL-1586^™^ ATCC), and human embryonic lung fibroblasts (HEL) cells (isolated at Amsterdam UMC) were used. HT-29, RD, Vero, and HEL cells were cultured in Eagle’s minimum essential medium (EMEM) (Lonza) supplemented with 8% heat-inactivated fetal bovine serum (FBS, Sigma-Aldrich), 100 U/mL penicillin/streptomycin (Pen-Strep) (Lonza), 0.1% (v/v) L-glutamine (Lonza), and 1% (v/v) non-essential amino acids (100x) (ScienceCell Research Laboratories). Vero E6 cells were cultured in Dulbecco’s Modified Eagle Medium (DMEM) (Gibco) supplemented with 10% FBS (Sigma-Aldrich), penicillin (100 U/ml; Lonza), and streptomycin (100 U/ml; Lonza). Cell lines were routinely cultured at 37°C with 5% CO2, 95% humidity and passaged every 7 days using 0.05% (v/v) Trypsin/EDTA (Gibco).

EV-A71 C1-91-480 was obtained from the RIVM (GenBank: AB552982.1) and was propagated in RD cells. HCMV isolate 3273 was derived from patient urine material and cultured on HEL fibroblasts. IAV isolate 61818 was derived from patient material and cultured on RD cells. PeV-A1 Harris strain was obtained from the RIVM and cultured on HT-29 cells. RV-A16 was derived from patient material and cultured on HEL cells. SARS-CoV-2 BetaCoV/Munich/BavPat1/2020 was obtained from Erasmus MC and cultured on Vero E6 cells. ZIKV strain H/PF/2013 was obtained from the Aix Marseille Université (Baronti et al., 2014) was cultured on Vero cells.

### 2.3 Human fetal airway cultures

#### 2.3.1 Isolation of primary airway epithelial cells

Airway epithelial cells were isolated from the bronchi and trachea of fetal donors (gestational age between 19 and 20 weeks and unknown gender). The isolation was performed as previously published (Fulcher et al., 2005). Cells were routinely cultured on rat tail collagen type I (Ibidi) coated T75 culture flasks in PneumaCult^TM^-Ex Plus medium (STEMCELL^TM^ Technologies) supplemented with 1% (v/v) Pen-Strep.

#### 2.3.2 Human fetal airway epithelium (HAE) cultures

Once primary airway epithelial cells reached ∼80% confluence, they were cultured on collagen coated cell culture inserts (Transwell^®^, 24-Well, 0.4 μm PET clear) (VWR) as previously published (Moreni *et al*., 2023). Cells were cultured in PneumaCult^TM^-Ex Plus medium supplemented with 10 μM Rock inhibitor for three days and without Rock inhibitor thereafter.

Once cells formed a confluent monolayer (2-5 days), cultures were differentiated by removing medium of the apical compartment and changing the basolateral cell culture medium to PneumaCult™-ALI medium (STEMCELL^TM^ Technologies) supplemented with 1% (v/v) Pen-Strep. Culture medium in the basolateral compartment was refreshed every 2–3 days. HAE cultures were maintained for at least 28 days in PneumaCult™-ALI medium before use. After 28 days of cultures, monolayers with transepithelial electrical resistance (TEER) >200 Ω x cm^2^ were used for further experiments.

### 2.4 Human intestinal enteroid cultures

#### 2.4.1 Isolation and maintenance of human intestinal enteroids

For the generation of fetal intestinal organoids, crypts were isolated from fetal intestinal tissue (gestational age between 16 and 18 weeks and unknown gender) as described previously (Sato et al., 2009). Isolated crypts were suspended in Matrigel^®^, dispensed in three 10 μL droplets per well in a 24-well tissue culture plate, and covered with 500 μL IntestiCult™ Organoid Growth Medium (STEMCELL^TM^ Technologies) supplemented with 1% (v/v) Pen-Strep. Enteroid cultures were maintained at 37°C, 5% CO2, medium was replenished every second day, and organoids were passaged every 3-5 days as described previously (Roodsant et al., 2020).

#### 2.4.2 Human fetal intestinal epithelium (HIE)

Cell culture inserts (cellQART^®^ 24-Well Insert 0.4 μm PET clear) were functionalized with human laminin (LN) suspension containing 10 μg/cm^2^ LN511 (BioLamina) and 5 μg/cm^2^ LN521 (BioLamina), for 2h at 37°C. After incubation, the LN suspension was removed. Human fetal enteroids were collected, and a single cell suspension was obtained by treatment with TrypLE (Gibco) for 10 min at 37°C. Cells were diluted to 5x10^5^ cells/mL in complete IntestiCult™ Organoid Differentation Medium (ODMh) (STEMCELL^TM^ Technologies) supplemented with 10 µM Rock inhibitor and 1% (v/v) Pen-Strep, and 200 μL of cell suspension was seeded per insert. A volume of 600 μL ODMh was added to the basolateral compartment. After three days in culture, the medium was changed to ODMh with 1% (v/v) Pen-Strep and refreshed every second day. On day 7, the cultures were placed on an orbital shaker at 66 rpm, medium was refreshed every second day for the basolateral compartment and daily for the apical compartment. After 14 days of culture, monolayers with TEER >200 Ω x cm^2^ were used for further experiments.

### 2.5 Transepithelial electrical resistance (TEER) measurements in HIE and HAE cultures

The TEER of the cell cultures seeded on inserts was measured using the epithelial volt-ohm meter EVOM2 Epithelial Volt ohmmeter (World precision instruments), routinely calibrated with a set of standard resistors. Resistance values of a coated empty insert were used as background. All TEER measurements were performed in triplicate per donor and the average was used for calculations. TEER values of >200 Ω x cm^2^ were considered adequate.

### 2.6 Stem cell culture and regionalized neural organoid (RNO) generation

Human pluripotent stem cells (hPSCs) lines, IMR90 (IMR90-4, WiCell), H1 (WAe001-A, WiCell) and Gibco (Gibco® Episomal hiPSC Line, Gibco^®^) were routinely cultured on treated 6-well tissue culture plates (CoStar^®^) maintained in mTeSR™ Plus (STEMCELL Technologies^TM^). Tissue culture plates were functionalised with 5 µg/mL LN521. Medium was refreshed daily, and stem cell lines were passaged weekly.

Dorsal forebrain regionalized neural organoids (RNOs) were generated using the STEMDiff™ Dorsal Forebrain Organoid Differentiation Kit (STEMCELL Technologies™) and STEMDiff™ Neural Organoid Maintenance Kit (STEMCELL Technologies™) according to the manufacturer’s protocols with minor modifications. Briefly, embryoid bodies (EB) were formed by Accutase™ (STEMCELL Technologies™) dissociation of hiPSCs and seeding 3.6 x10^6^ cells/well in AggreWell™800 plates (STEMCELL Technologies™). Medium was refreshed daily by removing and adding 1.5 mL basal medium. After six days, EBs were flushed from their AggreWell™ and transferred into pre-treated 6-mm tissue culture dishes and cultured on an orbital shaker (64 rpm). Tissue culture dishes were treated with Anti-Adherence Rinsing Solution (STEMCELL Technologies™). EBs were cultured in expansion medium until day 25 with medium changes every 2-3 days. On day 26, medium was changed to differentiation medium with medium changes every 2-3 days. On day 42, differentiation medium was changed to maintenance medium until the organoids were used for further experiments.

### 2.7 Viral infection and HH treatment

Lyophilized HH (10 mg) (Sigma-Aldrich) was reconstituted in dimethyl sulfoxide (DMSO, Sigma-Aldrich). HH was administered to HAE cultures at 40, 75 and 100 nM prepared in PneumaCult™-ALI medium without heparin. For HIE and RNOs, HH was administered at 4, 7.5 and 10 nM prepared in either ODMh or STEMDiff™ medium. All HH dosages were administered once at 0 hpi except for HCMV infected RNOs for which administration was performed every collection day. Dosages administered to HAE cultures were based on the intravenous biodistribution data of HH in CD2F1 mice and Ficher 344 rats. HH was found to accumulate within the lungs at a concentration of 160 ng/g (∼323 nM) and 98 ng/g (∼198 nM) at 48 and 72 hours respectively (Stecklair et al., 2001). Dosages administered to HIE and RNO cultures were based on the oral maximum plasma concentration Cmax that is achieved in humans. As part of a phase II clinical trial for the treatment of scleroderma, patients were treated with 3.5 mg/day HH, reaching a Cmax of 3.09 ng/mL or 6.235 nM (de Jonge et al., 2006). With Cmax values reported as ng/mL the micromolar concentration of a drug can be calculated using the formula below (Liston and Davis, 2017).

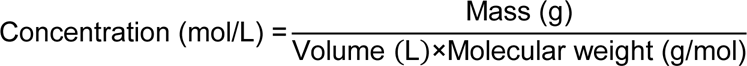

Viral infections on HIE and HAE were performed as described previously (García-Rodríguez et al., 2021; Karelehto et al., 2018; Sridhar et al., 2022). Briefly, viral stocks were diluted to 10^3.8^ TCID50/50 μL on ODMh for HIE or PneumaCult™-ALI medium without heparin for HAE. For apical infection, medium was removed from the apical side and 50 μL of viral inoculum was added. For basolateral infection, 50 μL of medium was removed and 50 μL of viral inoculum was added. See Table for the side of infection of the different viruses. The cultures were incubated for 2 h at 37°C with 5% CO2 and subsequently washed three times with Advanced Dulbecco’s Modified Eagle Medium/Nutrient Mixture F-12 (DMEM/F12 (Gibco)), both in the apical and the basolateral compartments. Medium containing specific concentrations of HH (0 µM, 4 µM, 7.5 µM, and 10 µM for HIE; and 0 µM, 40 µM, 75 µM, and 100 µM for HAE) was added to the cultures. Inserts were incubated for 10 min, after which the 0 hours post infection (hpi) time point was collected by removing 100 μL from the apical or the basolateral compartment. After collection, new medium was added (with or without HH) to both compartments for HIE and only to the basolateral compartment for HAE. Cultures were incubated for 72h, at which point the 72hpi was collected as described previously.

**Table 2.** List of viruses used to infect HAE and HIE models with their site of infection.

| **Virus** | **Model** | **Side of infection** | **Reference** |
| --- | --- | --- | --- |
| EV-A71 | HIE | Basolateral | (Aknouch et al., 2023) |
| PeV-A1 | HIE | Basolateral | (García-Rodríguez et al., 2021) |
| EV-A71 | HAE | Apical | (Hu et al., 2018) |
| RV-A16 | HAE | Apical | (Chen et al., 2006) |
| IAV | HAE | Apical | (Thompson et al., 2006) |
| SARS-CoV-2 | HAE | Apical | (Stölting et al., 2022) |

For the infection or RNOs, 48 days old organoids were used, except for ZIKV for which 28 days old organoids were used based on previous publications (Watanabe et al., 2017). For viral infections (except for HCMV), individual organoids were placed on a round bottom 96- well plate coated with Anti-Adherence Rinsing Solution and 100 μL of viral inoculum was added to each organoid. For ZIKV infection at 10^4.16^TCID50/50 μL was used and for EV-A71 and PeV-A1 infection 10^5^TCID50/50 μL was used. RNOs were incubated for 2 h at 37°C with 5% CO2 and washed three times with DMEM/F12 and moved to an anti-adherence coated 48-well plate containing 500 μL maintenance medium for EV-A71 and PeV-A1, and expansion medium for ZIKV. After 10 min incubation the 0hpi time point was collected by removing 180 μL medium, and fresh medium containing the appropriate concentrations of HH (0 µM, 4 µM, 7.5 µM, and 10 µM for EV-A71 and PeV-A1; and 0 µM, 1.3 µM, 4 µM, 7.5 µM for ZIKV) was added. RNOs were incubated for 72h, at which point the 72hpi was collected as described previously.

For HCMV infection, a different protocol was used due to its long replication cycle. RNOs were placed in an anti-adherence coated 48-well. Per well, medium was removed and 500 µL HCMV was added to the organoids at 10^2.9^TCID50/50 μL. RNOs were incubated with HCMV for six hours after which they were washed three times with DMEM/F12. After three washes, DMEM/F12 was replaced with maintenance medium containing 0 µM, 4 µM, 7.5 µM, and 10 µM HH. Medium containing the adequate concentrations of HH was refreshed on 2, 7, 12, and 17 dpi; at 21 dpi organoids were collected for DNA isolation.

### 2.8 Viability

#### 2.8.1 Intracellular ATP

Cell viability and cellular metabolic activity was determined based on intracellular adenosine triphosphate (ATP) production, using the 3D CellTiter-Glo Luminescent cell viability assay (Promega) as published previously (Wrzesinski et al., 2021) with minor modifications. Briefly, medium was aspirated and 100 µL PBS was added to each sample well of either HIE, HAE or RNOs. Cells were lysed with 100 µL 3D CellTiter-Glo lysis buffer and shaken in the dark for 30 min. Following incubation, the lysate was transferred to black clear bottom 96-well plates and the luminescence was measured in an H1 Synergy plate reader (BioTek). The data was normalized with reference to a standard curve for ATP (Sigma-Aldrich) and relevant controls.

#### 2.8.2 Extracellular LDH

Cell toxicity was determined by extracellular lactate dehydrogenase (LDH) release, using the LDH-Glo™ Cytotoxicity assay (Promega) following manufacturer’s instructions. Briefly, culture medium samples (5 µL) were added to 95 µL LDH storage buffer and stored at -70°C until further processing. The assay was performed by transferring 50 µL of the sample and LDH standard in LDH storage buffer to a black clear bottom 96-well plate. LDH detection reagent (50 µL) was added to standard and sample wells and incubated at room temperature in the dark for one hour. Luminescence was measured in an H1 Synergy plate reader (BioTek). Data was normalized with reference to a standard curve of LDH and relevant controls.

### 2.9 Quantitative RT-qPCR

#### 2.9.1 RNA isolation and reverse-transcription

RNA was isolated from 25 μL of collected supernatant of infected cultures using the PureLink^TM^ RNA Mini kit (Thermo Fisher) according to the manufacturer’s instructions. Equal volumes of eluted RNA were used for reverse-transcription using the SuperScript™ II Reverse Transcriptase synthesis kit (Thermo Fisher). For SARS-CoV-2, RNA isolation and reverse-transcription was performed as described previously (Aknouch et al., 2022).

#### 2.9.2 DNA isolation

For HCMV, whole RNOs were collected at 21 dpi and viral DNA was isolated using the ISOLATE II Genomic DNA Kit (Meridian Bioscience) according to the manufacturer’s instructions.

#### 2.9.3 qPCR

A volume of 5 μL of cDNA was used for quantitative PCR (qPCR) on a CFX Connect Real-Time PCR Detection System (Bio-Rad). TaqMan and SYBR Green qPCRs were performed as described previously (Benschop et al., 2010). For SARS-CoV-2 RT-qPCR was performed as described previously (Aknouch et al., 2022). Cq values were transformed into viral genome copies using a standard curve for each specific set of primers, see Supplementary Table 2. In the case of HCMV, genome copies were also normalized to the RNO’s size based on the expression of β-actin.

#### 2.10 Median tissue culture infectious dose (TCID50)

Supernatant samples at 0 and 72hpi were used to determine the presence of infectious viral particles, titrations were performed on the cell lines used to make stocks (see above). Seven ten-fold dilutions of each sample were performed and 50 µL of each dilution was added to a 96-well plate, after which 200 µL of specific cells were added. Plates were incubated for seven days until scoring of the cytopathic effect (CPE) and calculation of the TCID50 according to the Reed and Muench Method (Reed and Muench, 1938). Values at 72hpi were normalized to 0hpi to determine the increase of infectious particles.

#### 2.11 Proline content determination in viruses and host factors

Proline content and presence of PP-motifs were quantified for the viruses used in this study and the important host factors for their infection, such as receptors. Protein sequences were obtained from UniProt database and quantification was performed as proline percentage of the total amount of amino acids per protein.

2.12 Statistics

Unless otherwise stated, HAE and HIE experiments were conducted using three independent donors in technical duplicates, while RNOs experiments were conducted with three different iPSCs cell lines and technical triplicates. Data are expressed as mean ± standard error of the mean (SEM), unless otherwise indicated. Statistical analysis was conducted using GraphPad Prism 8 software (GraphPad Software Inc.). The specific statistical tests performed for each analysis are stated in the corresponding figure legend. Statistical significance was accepted at *p*-value < 0.05.

## 3 Results

### 3.1 Halofuginone hydrobromide at clinically relevant concentrations does not affect organoid viability

To determine the effect of the chosen HH concentrations, based on biodistribution and Cmax values, on HAE, HIE and RNO cultures, viability and metabolic activity were analyzed by measuring the intracellular ATP levels at 72 hours post treatment and extracellular LDH release, as an indication of cell toxicity, was measured at 24-hour intervals. For the HAE cultures, the intracellular ATP concentration normalized to the untreated control remained ≥100% for all the administered HH concentrations (Figure 2A). The limited cellular toxicity was confirmed by the low extracellular LDH release for all concentrations over time (Figure 2B). Similarly, the intracellular ATP concentration for the HIE cultures remained above 95% for all HH concentrations (Figure 2C), while also showing low LDH release (Figure 2D), except for the 0hpi time-point, with higher release probably due to the manipulation of the cultures. RNOs showed slight sensitivity to HH at 4 and 7.5 nM with viability measured at ≥80% while treatment with a 10 nM HH concentration resulted in viability of ≥90% following a 72-hour one-time administration (Figure 2E). We did not detect a significant increase in extracellular LDH release when compared to the untreated control, indicating no cell toxicity upon treatment with all HH concentrations for 72 hours (Figure 2F). Moreover, RNOs treated with HH at 4, 7.5 and 10 nM for 21 days showed only a non-significant increase in extracellular LDH release for HH at 4 nM on day 7, whereas for days 12, 17 and 21 LDH release for all concentrations were below that of the untreated control (Supplementary Figure 1).

**Figure 2.**
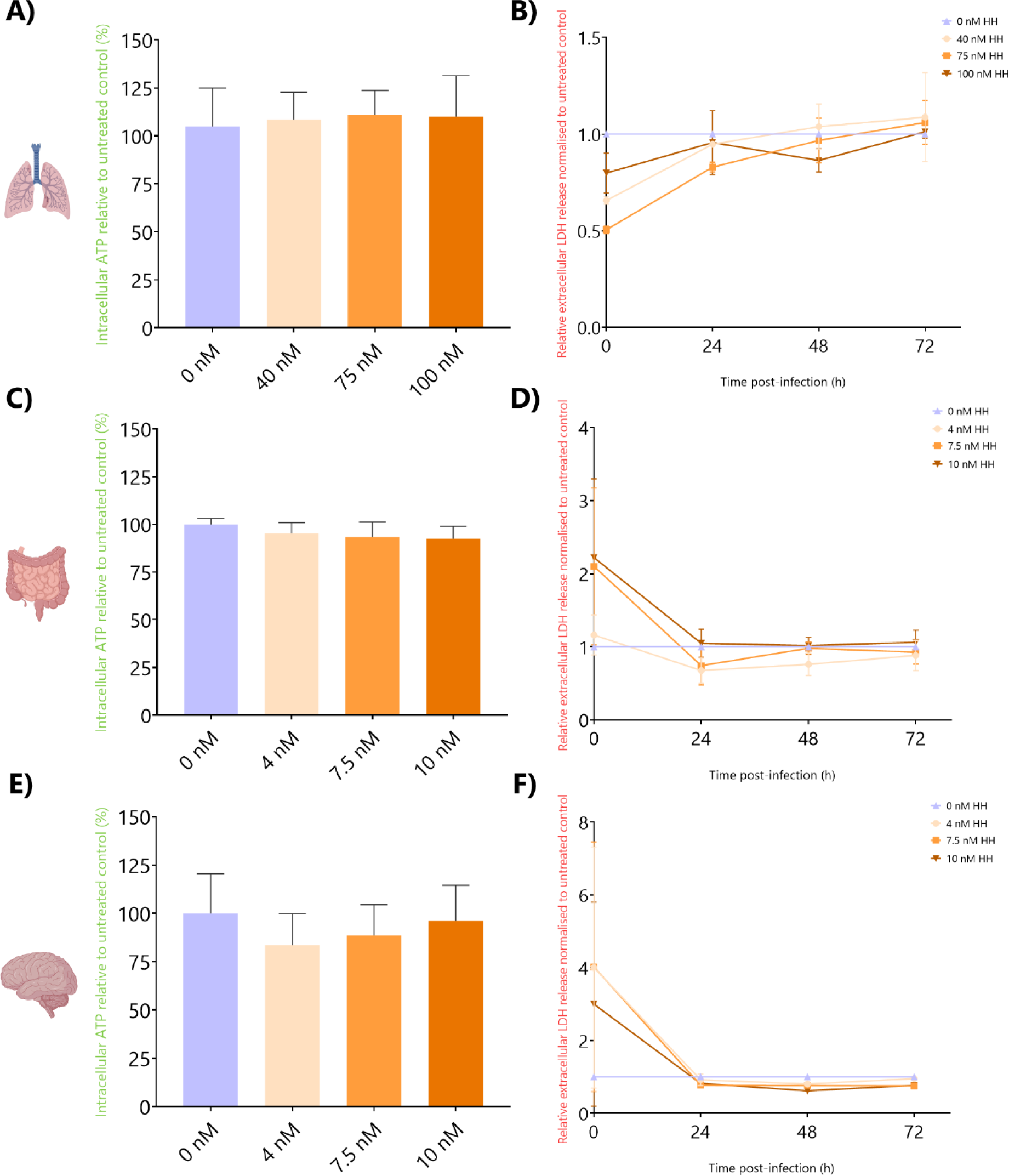
Organoid viability after treatment with one dose of HH at different concentrations. Intracellular ATP relative to untreated controls (%) in HAE **(A)**, HIE **(C)** and RNOs **(E)** treated with different concentrations of HH. Relative LDH release normalized to untreated control in HAE **(B)**, HIE **(D)** and RNOs **(F)**. In all cases, data represents the mean ± SEM of HAE/ HIE three biological replicates in technical duplicates, and RNOs three biological replicates in technical triplicates.

### 3.2 Halofuginone hydrobromide inhibits SARS-CoV-2 replication in HAE

We determined the effect of HH on HAE infected with a panel of four viruses, EV-A71, IAV, RV-A16 and SARS-CoV-2. HH treatment of HAE infected with EV-A71 or IAV did not lead to a significant reduction in the viral copy numbers at any of the tested concentrations (Figure 3A and D). In line, we observed no significant alterations in the infectious viral particles, as assessed by TCID50, regardless of the HH dosage (Figure 3B and E). We did not observe any significant changes in metabolic activity (Figure 3A and D) or extracellular LDH release (Figure 3C and F) for any of the tested concentrations. HAE cultures infected with RV-A16 and treated with HH did not show a significant decrease in viral RNA copies (Figure 3G) nor in the amount of infectious RV-16 particles for all HH concentrations (Figure 3H). The metabolic activity of the HAE cultures (Figure 3G) and extracellular LDH release is maintained between the conditions (Figure 3I). HH treated HAE cultures infected with SARS-CoV-2 showed a concentration dependent decrease in viral copy numbers and infectious viral particles at 75 and 100 nM (Figure 3J and Figure 3K respectively). Significant decrease in the metabolic activity of the HAE cultures at 72h was observed for the highest HH concentration (Figure 3J) but no significant increase in extracellular LDH release was observed (Figure 3L).

**Figure 3.**
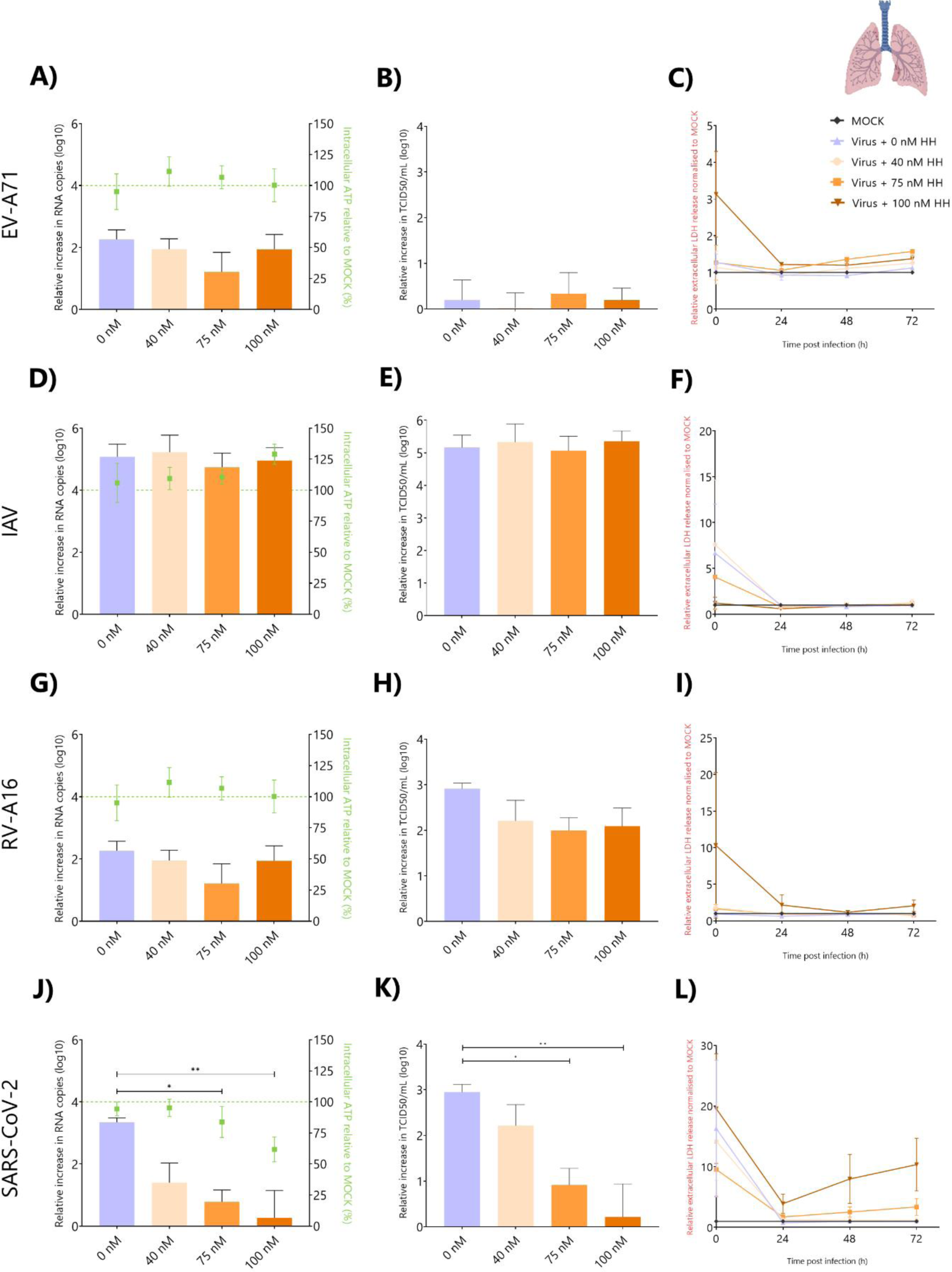
HH effect on viral infection on HAE cultures. **(A, D, G, F)** Relative increase in viral genome copies at 72hpi treated with different concentrations of HH and the corresponding intracellular ATP relative to MOCK infected cultures. Values below the dashed green line represent lower intracellular ATP relative to MOCK infected cultures **(B, E, H, K)** Relative increase in virus titer at 72hpi in untreated or HH treated HAE cultures. **(C, F, I, L)** Relative LDH release of HAE infected cultures and treated with different concentrations of HH as compared to MOCK infected cultures at 0, 24, 48, and 72 hpi. **(A-C)** Correspond to EV-A71 infected HAE cultures, **(D-F)** to IAV infected HAE cultures, **(G-I)** to RV-A16 infected HAE cultures, and **(J-L)** to SARS-CoV-2 infected HAE cultures. In all cases, data represents the mean ± SEM of three biological replicates in technical duplicates. Statistical significance was analysed with One-way ANOVA with Tukey’s multiple comparisons. * p-value < 0.05; ** p-value <0.01.

### 3.3 Halofuginone hydrobromide does not inhibit EV-A71 or PeV-A1 replication in HIE

HIE cultures were infected with EV-A71 and PeV-A1 and treated with HH at 4, 7.5 and 10 nM. Treatment of EV-A71 or PeV-A1 infected HIE cultures with any concentration of HH did not reduce viral copy numbers or infectious viral particles as compared to the untreated EV-A71 and PeV-A1 conditions (Figure 4A and Figure 4D respectively). For both EV-A71 and PeV-A1 infected HIE cultures, we noticed a decrease of ≥25% overall metabolic activity and viability for all experimental groups (Figure 4A and Figure 4D respectively) that were significantly lower than the MOCK (Supplementary Figure 2E and Supplementary Figure 2F respectively). The decrease in cell metabolic activity and viability correlated to an increase in extracellular LDH at 24 hpi for all conditions of both EV-A71 and PeV-A1 infected cultures when compared to the untreated control (Figure 4C and Figure 4F respectively).

**Figure 4.**
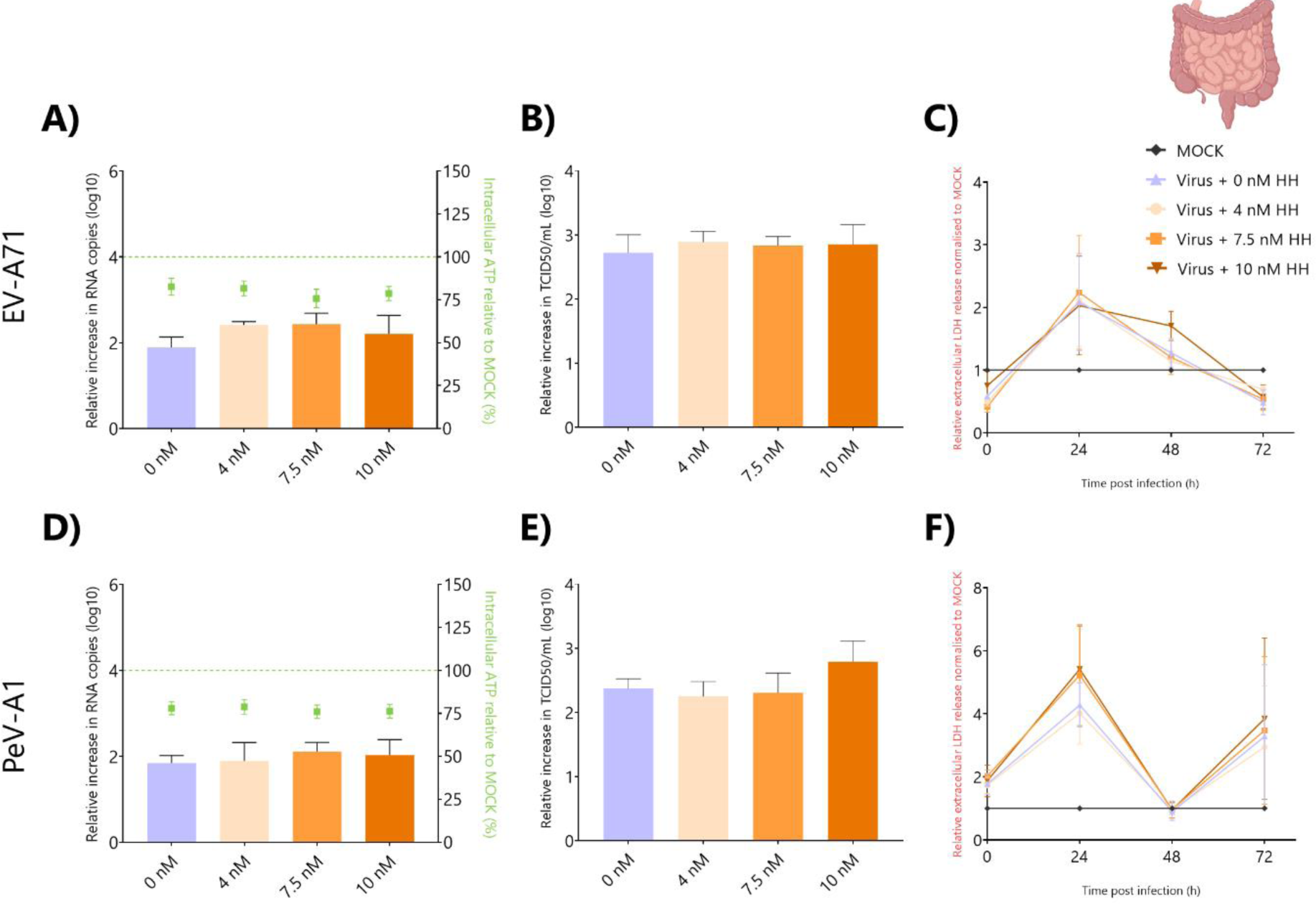
HH effect on viral infection on HIE cultures. **(A, D)** Relative increase in viral genome copies at 72hpi treated with different concentrations of HH and the correspondent intracellular ATP relative to MOCK infected cultures. Values below the dashed green line represent lower intracellular ATP relative to MOCK infected cultures (**B, E)** Relative increase in virus titer at 72hpi in untreated or HH treated HIE cultures. **(C)** Relative LDH release of HIE infected cultures and treated with different concentrations of HH as compared to MOCK infected cultures at 0, 24, 48, and 72 hpi. **(A-C)** Correspond to EV-A71 infected HIE cultures, **(D-F)** to PeV-A1 infected HIE cultures. In all cases, data represents the mean ± SEM of three biological replicates in technical duplicates.

### 3.4 Halofuginone hydrobromide does not inhibit EV-A71, PeV-A1, HCMV or ZIKV replication in RNOs

RNOs infected with EV-A71 and PeV-A1 and treated with HH did not show a significant reduction in the viral copy numbers or infectious viral particles (Figure 5A-B and Figure 5D-E, respectively). EV-A71 infected RNOs, showed a decrease of ≥10% in metabolic activity (Figure 5A), while PeV-A1 infected cultures showed a decrease of ≥25% for all experimental groups (Figure 5D). For EV-A71 infected cultures, we observed no alterations in extracellular LDH at 24 and 72 hpi (Figure 5C), while LDH release for PeV-A1 infected RNOs was reduced when compared to the untreated MOCK at 24 and 48hpi (Figure 5F). HCMV infected RNOs treated with HH did not show any alterations in viral copy numbers and had a ≥25% decrease in metabolic activity and viability for all experimental groups (Figure 5G). The loss in viability and cellular metabolic activity correlated to the observed increase in extracellular LDH release at 2 and 7 dpi (Figure 5H). Similarly, no significant decrease in viral copy numbers was observed in ZIKV infected RNOs following HH treatment (Figure 5I). However, toxicity caused by ZIKV infection is noted in significant decreases in viability and cellular metabolic activity of ≥50% for all experimental groups at 72 hpi (Figure 5I and Supplementary Figure 2J).

**Figure 5.**
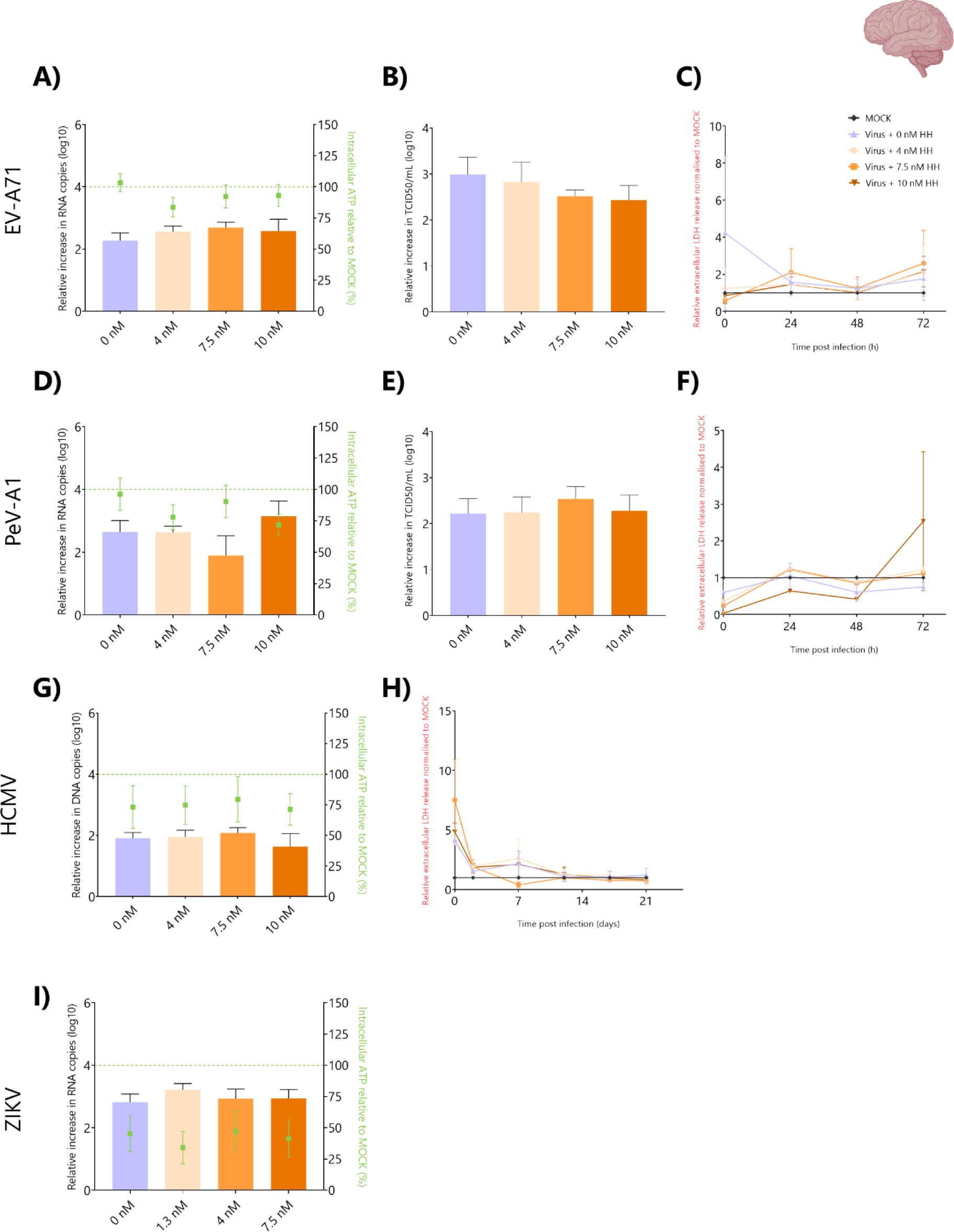
HH effect on viral infection on RNOs. **(A, D, G, I)** Relative increase in viral genome copies at 72hpi or 21dpi **(G)** treated with different concentrations of HH and the correspondent intracellular ATP relative to MOCK infected cultures. Values below the dashed green line represent lower intracellular ATP relative to MOCK infected cultures (**B, E)** Relative increase in virus titer at 72hpi in untreated or HH treated RNOs. **(C, F, H)** Relative LDH release of RNOs infected and treated with different concentrations of HH as compared to MOCK infected cultures at 0, 24, 48, and 72 hpi. **(A, F)** or 7, 14, 21 dpi **(H)**. **(A-C)** Correspond to EV-A71 infected RNOs, **(D-F)** to PeV-A1 infected RNOs, **(G-H)** to HCMV infected RNOs, and **(I)** to ZIKV infected RNOs. In all cases, data represents the mean ± SEM of three biological replicates in technical triplicates, except for HCMV and PeV-A1 for which two biological replicates were used.

### 3.5 Proline content and PP-motifs are similar within viruses and host factors

As HH binding to prolyl tRNA synthetase leads to an accumulation of uncharged prolyl tRNAs, proline content could have an effect on the antiviral activity of HH. Proline content was similar for most of the host factors important for viral infection of the viruses used in this study, with intercellular adhesion molecule 1 (ICAM-1), one of the receptors for RV-A16, having a total proline content of 9.4%, with 13.8% in the topological cytoplasmic domain of the protein. Interestingly, when focusing on specific protein regions, we found that the transmembrane serine protease 2 (TMPRSS2), an important host factor for SARS-CoV-2 has an abundance of proline (19.1%) within its topological cytoplasmic domain. Furthermore, although angiotensin-converting enzyme 2 (ACE2), the main receptor for SARS-CoV-2, does not have an abundance of proline, 4.6%, it does have proline in a regions important for viral binding. Additionally, PP-motifs were found to be uniform for the host factors described (Supplementary Table 4). Along the same lines, the proline content of the viruses used in this study was similar, between 3.8% and 5.9%, with HCMV showing the highest content. In contrast, PP-motifs were not equally present in all the viruses, for instance PeV-A1 only has one PP-motif, while HCMV has several (Supplementary Table 5).

## 4 Discussion

For the current study, we assessed the toxicity and antiviral potential of HH at three concentrations for 72 hours against a panel of viruses (EV-A71, PeV-A1, IAV, RV-A16, HCMV, SARS-CoV-2 and ZIKV) in three different organoid models (HAE, HIE and RNO). Furthermore, we also assessed the proline content and presence of PP-motifs for both the evaluated viruses and the known viral host factors.

It is noteworthy that during pre-clinical *in vitro* studies drugs are administered at concentrations that far exceed concentrations that can be achieved in a clinical setting (Liston and Davis, 2017). This approach renders these *in vitro* studies ineffective at providing clinically relevant and translatable data. We therefore aimed to increase the predictive value of antiviral effectiveness of HH for the clinical setting by two methods. Firstly, by calculating clinically relevant HH concentrations corresponding to available *in vivo* pharmacokinetic biodistribution or Cmax data of HH. Secondly, by utilizing HAE, HIE and RNO organoid models representing the human primary and secondary replication site of the evaluated virus panel. With this in consideration, our results indicate that HH at clinically relevant concentrations does not significantly affect the viability of our HAE and HIE following a 72-hour exposure, while similar observations were made for RNOs following 72-hour and 21-day HH exposure. During the assessment of HH efficacy we found that, similar to Chen *et al* 2021, our results indicated that HH showed significant efficacy against SARS-CoV-2, however no efficacy was observed for

EV-A71, PeV-A1, IAV, RV-A16, HCMV, or ZIKV in human organoid models. The fact that HH did not show any efficacy against the other viruses was unexpected as the possible broad-spectrum antiviral efficacy of HH was indicated before (Chen et al., 2021). The lack of a broad-spectrum antiviral activity against these selected viruses could potentially be explained by examining its molecular target.

The molecular target of HH is Prolyl-tRNA synthetase 1 (PARS1) an aminoacyl-tRNA synthetase (ARSs), that facilitates the covalent ligation of proline to proline-tRNAs for protein synthesis (Hwang et al., 2019; Keller et al., 2012; Yoon et al., 2023). The four codons that code for proline is recognized by three distinct tRNAs, which are all charged by PARS1. Aside from proline-tRNAs being dependent on PARS1, proline, the only n-alkyl amino acid, has some unique chemical properties. This is seen in the effect that proline and its nascent peptide has on the translation and elongation speed of messenger RNA (mRNA). The speed of peptide bond formation is significantly influenced by polyproline (PP or PPP) motifs, with abundant PP-motifs resulting in ribosome pauses that significantly slows protein translation. It has been previously stated that the efficacy of HH, a competitive inhibitor to the catalytic activity of PARS1, is specific to proline-rich proteins rather than proline rare proteins (Krafczyk et al., 2021; Yoon et al., 2023). However, the abundance of PP-motifs also plays a role and this is evident in the reduced biosynthesis of collagen I, a protein abundant in proline and PP-motifs, following HH treatment (Yoon et al., 2023), see Supplementary Table 3.

Following the evaluation of the proline and PP-motifs of the viruses and their respective host factors for this study, we found that ACE2 and TMPRSS2, the receptor and host factor for SARS-CoV-2, have prolines in important binding areas and an abundance of proline in specific protein regions. These include the substrate binding region (345-346, HP) as well as the so called MYP region coded by aa 82-84. Mutagenesis in this MYP region on the ACE2 receptor significantly inhibited interaction with the SARS-CoV-1 spike glycoprotein that may hold significance for SARS-CoV-2 (Li et al., 2005). We further expected HH to also show significant efficacy against RV-A16 and HCMV. This is due to the fact that the entry receptor for RV-A16, ICAM-1 has abundant proline and RV-A16 and HCMV have abundant PP-motifs. However, for RV-A16 we did not observe a decrease in viral copy numbers; nonetheless, we do note a decrease albeit not significant in the amount of infective virus particles of RV-A16 following treatment with HH. Evaluating the abundance of proline and PP-motifs in our virus panel (Supplementary Table 5), we found that RV-A16 peptide not only had the most abundant proline of all the viruses in our panel, also had various PP, PPP and PPPP motifs. Therefore, as HH directly influences translation it would explain why we noticed a decrease in infectious viral particles, with no effect on the viral copy numbers. Similarly, HCMV also has numerous PP, PPP and PPPP motifs. The lack of HH efficacy may be found in the high fidelity of protein translation that is governed by robust post-transcriptional regulation. HH treatment results in increase in uncharged prolyl-tRNAs causing AAR and ribosomal pauses. Acting as a ribosomal pause release factor during translation is the highly conserved, eukaryotic initiation factor 5A (eIF5A). One of its main functions is to facilitate the translation of PP-motifs (Schuller et al., 2017). Simultaneously, during viral infection, eIF5A is upregulated to promote viral translation (Seoane et al., 2022). The upregulation of eIF5A may counteract the effect of HH, and therefore not completely inhibit viral protein formation. However, further experiments are needed to confirm this hypothesis.

In the current study we also expected to see HH efficacy against ZIKV as previously shown. Additionally, it also shares important host factors with DENV and CHIKV for which HH antiviral activity have also been shown (Hwang et al., 2019). The reason for this discrepancy may be found in the concentrations of HH and the *in vitro* model used in previous reports. Hwang *et al*., 2019, showed HH efficacy against DENV and CHIKV following the infection of liver hepatoma (Huh-7) cells at 25, 50 and 100 nM. Although these concentrations are clinically relevant for the liver, they are not for the brain (Stecklair et al., 2001). Within *in vivo* models no HH reached or accumulated within the brain. HH (53 ng/g, ∼106 nM) was found within rat brain tissue at 3.0 mg/mL at 15 min, but significantly declined after 3hrs, with no HH detectable within the tissue thereafter (Stecklair et al., 2001). Thus, the differences in the concentration of HH administered within our RNO model could explain the lack of viral inhibition for ZIKV. It could also be argued that employing models not representative of the *in vivo* situation negates HH efficacy especially if the results are not translatable to the *in vivo* situation. The primary replication site for ZIKV has been identified as the skin, testis and placenta with the secondary infection site being embryonic brain tissue, not the liver (Supplementary Table 1). Again, highlighting the importance of models like human RNOs that are more physiologically relevant to provide clinically translatable data.

In summary, using known biodistribution and Cmax values of repurposed drug will allow us to provide more clinically relevant predictions. While organoid models provide the biomimetic setting to predict physiologically relevant and clinically translatable readouts during screening of antiviral efficacy. With focus on HH we suggest that the antiviral efficacy thereof is not only related to the concentration of HH administered, but also the proline content and the presence of PP-motifs in the host receptors.

## 5 Conclusion

Although HH has only shown efficacy against SARS-CoV-2 in this study it may still hold antiviral efficacy for other potential circulating or emerging human viruses. Although HH is suggested to be a broad-spectrum antiviral, we only found HH antiviral activity against SARS-CoV-2 but not against other viruses in physiologically relevant models and HH administered at physiologically relevant concentrations. However, it appears as though the efficacy of HH is not only concentration dependent but also upon the abundance of proline within the host factors and where within the protein this proline is found.

## 6 List of abbreviations

AAR: amino acid starvation response
ACE2: angiotensin-converting enzyme 2
ARS: aminoacyl-tRNA synthetase
ATP: adenosine triphosphate
CHIKV: chikungunya virus
CPE: cytopathic effect
DENV: dengue virus
DMEM/F12: Advanced Dulbecco’s Modified Eagle Medium/Nutrient Mixture F-12
DMEM: Dulbecco’s Modified Eagle Medium
EB: embryoid body
EMEM: Eagle’s minimum essential medium
EV-A71: enterovirus A71
FBS: fetal bovine serum
HAE: human airway epithelium
HCMV: human cytomegalovirus
HH: halofuginone hydrobromide
HIE: human intestinal epithelium
hpi: hours post infection
IAV: influenza A virus
ICAM-1: intercellular adhesion molecule 1
LDH: lactate dehydrogenase
LN: laminin
mRNA: messenger RNA
ODMh: IntestiCult™ Organoid Differentation Medium
PARS1: Prolyl-tRNA synthetase 1
Pen-Strep: penicillin/streptomycin
PeV-A1: parechovirus A1
qPCR: quantitative polymerase chain reaction
RNO: regionalized neural organoid
SARS-CoV-2: severe acute respiratory syndrome coronavirus 2
SEM: standard error of the mean
TEER: transepithelial electrical resistance
TMPRSS2: transmembrane serine protease 2
ZIKV: Zika virus

## Acknowledgements

All schematic representations were created with <u>BioRender.com</u>. Authors would like to acknowledge Hetty van Eijk for her work on the last publication of her career and wish she enjoys her retirement.

## 1. Supplementary data

**Supplementary Table 1.** Characteristics of the different viruses used in this study. GI: gastrointestinal tract. For genomes: +ssRNA (positive sense single stranded RNA genome), ddDNA (double stranded DNA genome), -ssRNA (negative sense single stranded RNA genome).

| Virus | Family and Genus | Genome | Enveloped/<br>Non-<br>enveloped | Clinical indications | Receptors | Primary<br>replication<br>site | Secondary<br>replication<br>site | Incidence | Ref |
| --- | --- | --- | --- | --- | --- | --- | --- | --- | --- |
| Enterovirus A71 (EV-A71) | *Picornaviridae*<br>*Enterovirus* | +ssRNA | Non<br>enveloped | Hand, foot, and mouth disease<br>herpangina, diarrhea,<br>aseptic<br>meningitis, acute flaccid<br>paralysis,<br>encephalitis, myocarditis,<br>delayed neurodevelopment,<br>reduced cognitive function. | Human scavenger<br>receptor class B<br>member 2 (SCARB2) | GI and upper<br>respiratory tract | Brain, heart | Several outbreaks<br>worldwide since<br>isolation in 1969 with<br>1,000-100,000 recorded<br>cases and mortality<br>between 50-1000 | (Li et al., 2021;<br>Puenpa et al.,<br>2019; Wang et al.,<br>2022; Yuan et al.,<br>2018) |
| Human cytomegalovirus<br>(HCMV) | *Herpesviridae*<br>*Cytomegalovirus* | ddDNA | Enveloped | Occasional cases of infectious<br>mononucleosis, end organ<br>disease in<br>immunocompromised patients,<br>congenital infection that can<br>lead to neurodevelopmental<br>abnormalities in infants and<br>children. | Platelet Derived<br>Growth Factor<br>Receptor Alpha<br>(PDGFRA) and<br>neuropilin2 (Nrp2) | GI tract, lungs,<br>eyes | Brain,<br>salivary glands,<br>placenta | Seroprevalence ranges<br>from 25% to 95%<br>globally | (Fowler et al.,<br>2022; Gerna et al.,<br>2019; Griffiths and<br>Reeves, 2021) |
| Influenza A virus (IAV) | *Orthomyxoviridae*<br>*Alphainfluenzavirus* | Segmented,<br>-ssRNA | Enveloped | Fever, headache, weakness,<br>red eyes, sore throat, dry<br>cough, and nasal discharge | Sialic acids | Upper<br>respiratory tract | The respiratory<br>epithelium is the only<br>site where the<br>hemagglutinin<br>molecule is<br>effectively cleaved,<br>generating infectious<br>virus particles | Globally 1 billion<br>infections, with 4 million<br>severe infection cases<br>and<br>500,000 deaths annually | (Javanian et al.,<br>2021; Kalil and<br>Thomas, 2019;<br>Krammer et al.,<br>2018) |
| Parechovirus A1 (PeV-A1) | *Picornaviridae*<br>*Parechovirus*, | +ssRNA | Non enveloped | Mild gastroenteritis and respiratory symptoms. Rare cases of acute flaccid paralysis and meningitis. | Myeloid-associated differentiation marker (MYADM), integrins. | GI and upper respiratory tract | In rare cases presence of the virus in the CSF | Seroprevalence ranges from 92% to 99% globally. | (Figueroa et al., 1989; Romero and Selvarangan, 2011; Sridhar et al., 2019; Watanabe et al., 2016) |
| Rhinovirus A16 (RV-A16) | *Picornaviridae*<br>*Enterovirus* | +ssRNA | Non enveloped | Rhinorrhea, nasal congestion, sore throat, cough, **headache, malaise acute, otitis media**, rhinosinusitis and croup | Intercellular adhesion molecule 1 (ICAM-1) and low density lipoprotein receptor (LDL-R) | Upper and lower respiratory tract | Infection is limited to the respiratory tract with only rare reports of invasive disease | Worldwide 11–20 million hospitalizations and 2 million deaths annually in children <5 years | (Annamalay et al., 2012; Bizot et al., 2021; Jacobs et al., 2013) |
| Severe acute respiratory syndrome coronavirus 2 (SARS-CoV-2) | *Coronaviridae*<br>*Betacoronavirus* | +ssRNA | Enveloped | Mild to moderate respiratory disease, experiencing cough, fever, headache, myalgia and diarrhea. Severe disease is dyspnea, progressive respiratory failure can occur in patients with severe COVID-19 | Angiotensin-converting enzyme 2 (ACE2) | Nasopharynx, trachea, nasal olfactory mucosa | SARS-CoV-2 RNA has been detected in several specimens, such as feces, blood or CSF. But it is unknown if this presence is due to direct infection. | Globally, ≥500 million confirmed cases of COVID-19 November 2021, over 3 billion individuals, or 44% of the world's population, had been infected with SARS-CoV-2 at least once | (Collaborators, 2022; Lamers and Haagmans, 2022) |
| Zika virus (ZIKV) | *Flaviviridae*<br>*Flavivirus* | +ssRNA | Enveloped | Fever, arthralgia, myalgia, conjunctivitis, headache, fatigue and/or **malaise, dizziness, stomachache**<br>**Rare symptoms:** multi-organ failure, meningitis and encephalitis, and thrombocytopenia | Tyrosine-protein kinase receptor UFO (AXL), dendritic cell-specific intercellular adhesion molecule-3-grabbing non-integrin (DC-SIGN), tyrosine-protein kinase receptor (TYRO3). | Skin, testis, placenta | Embryonic brain tissue | Cases of ZIKV per year: 81,852 (2016), 609 (2017), 1,800 (2018) and 15 cases (2019). ~440 000–1300 000 cases in Brazil during the 2016 outbreak | (Bhargavi and Moa, 2020; Pielnaa et al., 2020) |

**Supplementary Table 2.**
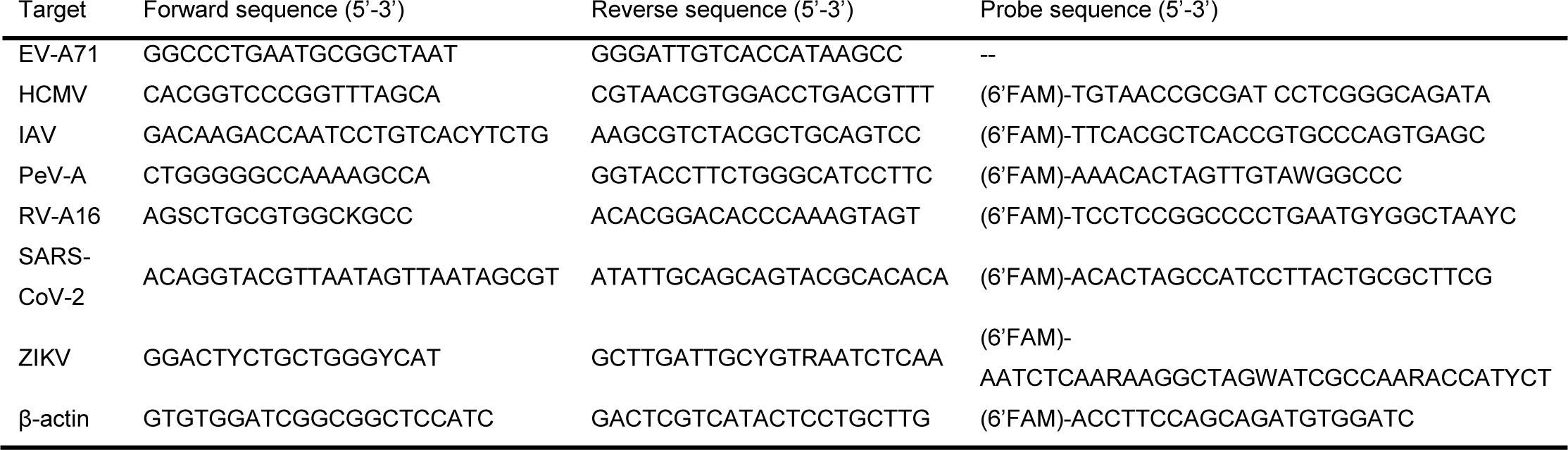
List of primers used in this study with forward and reverse sequences.

**Supplementary Table 3.**
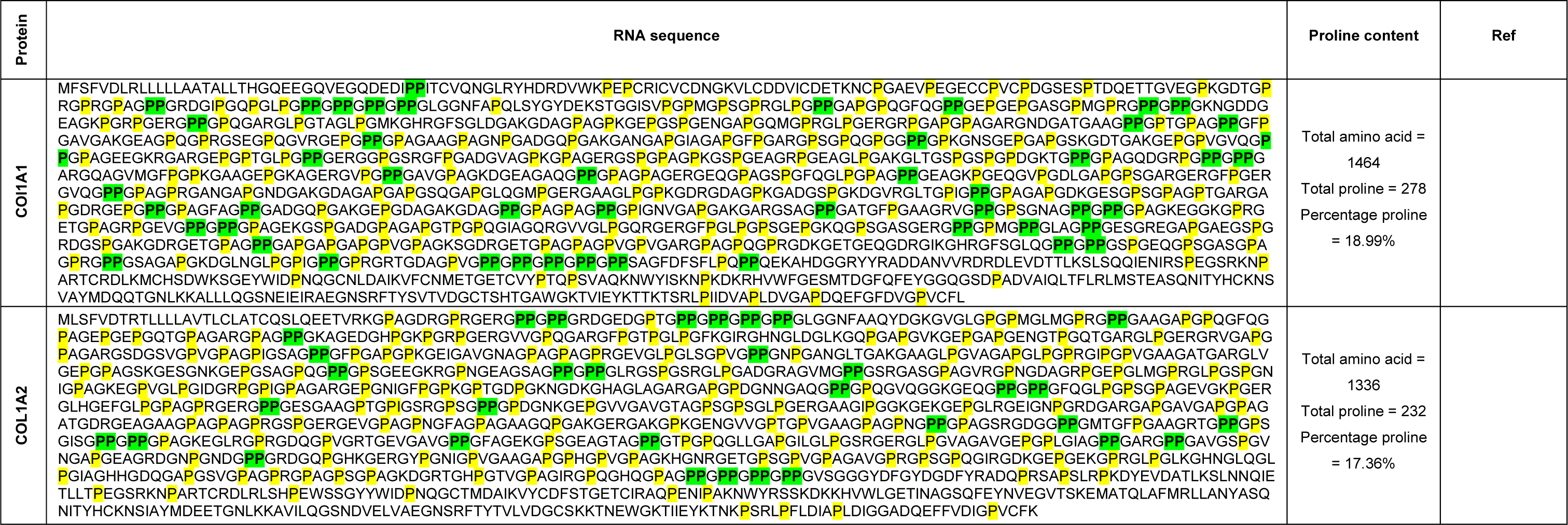
Estimated proline content and location of collagen.

**Supplementary Table 4.**
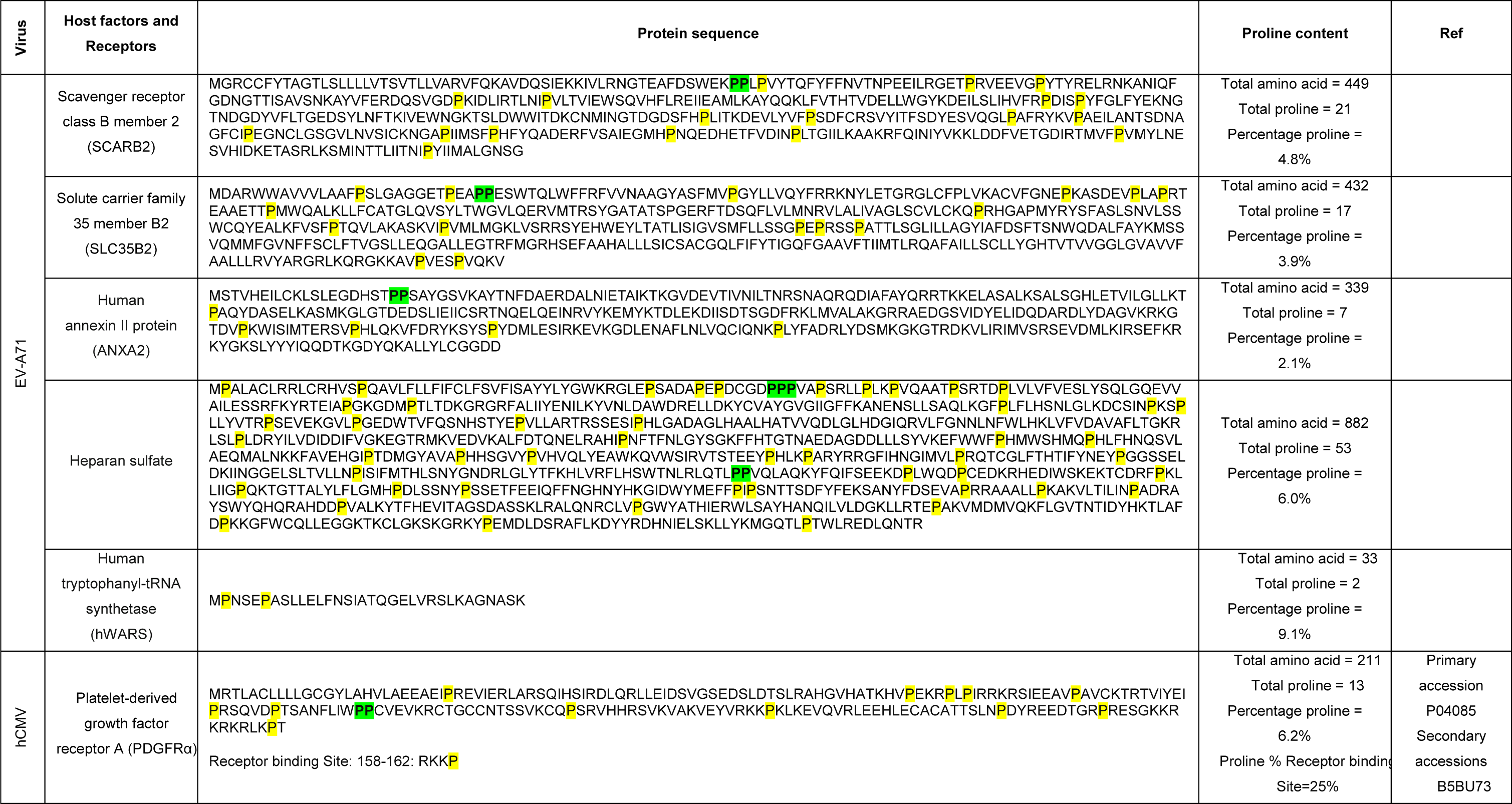
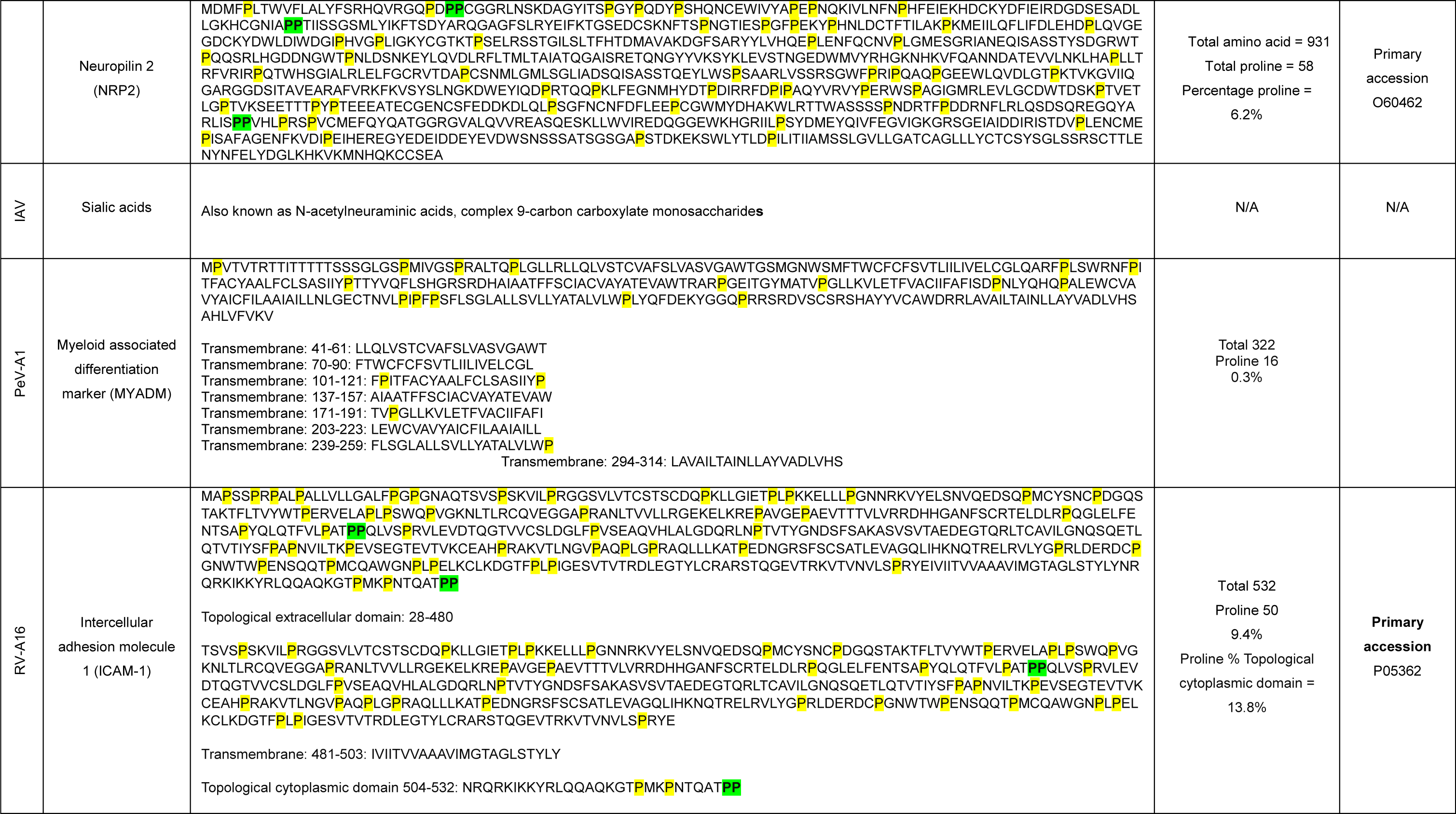
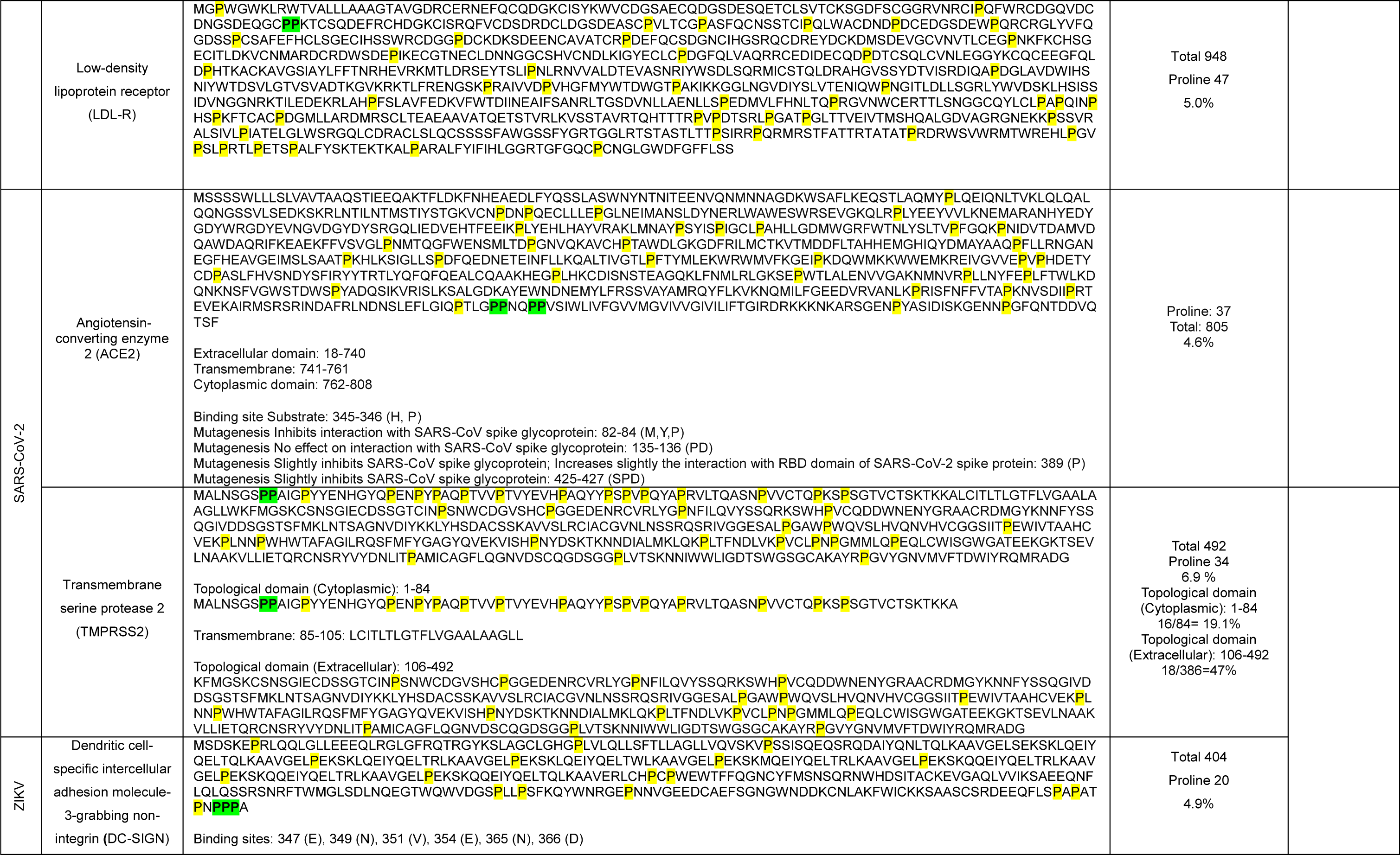
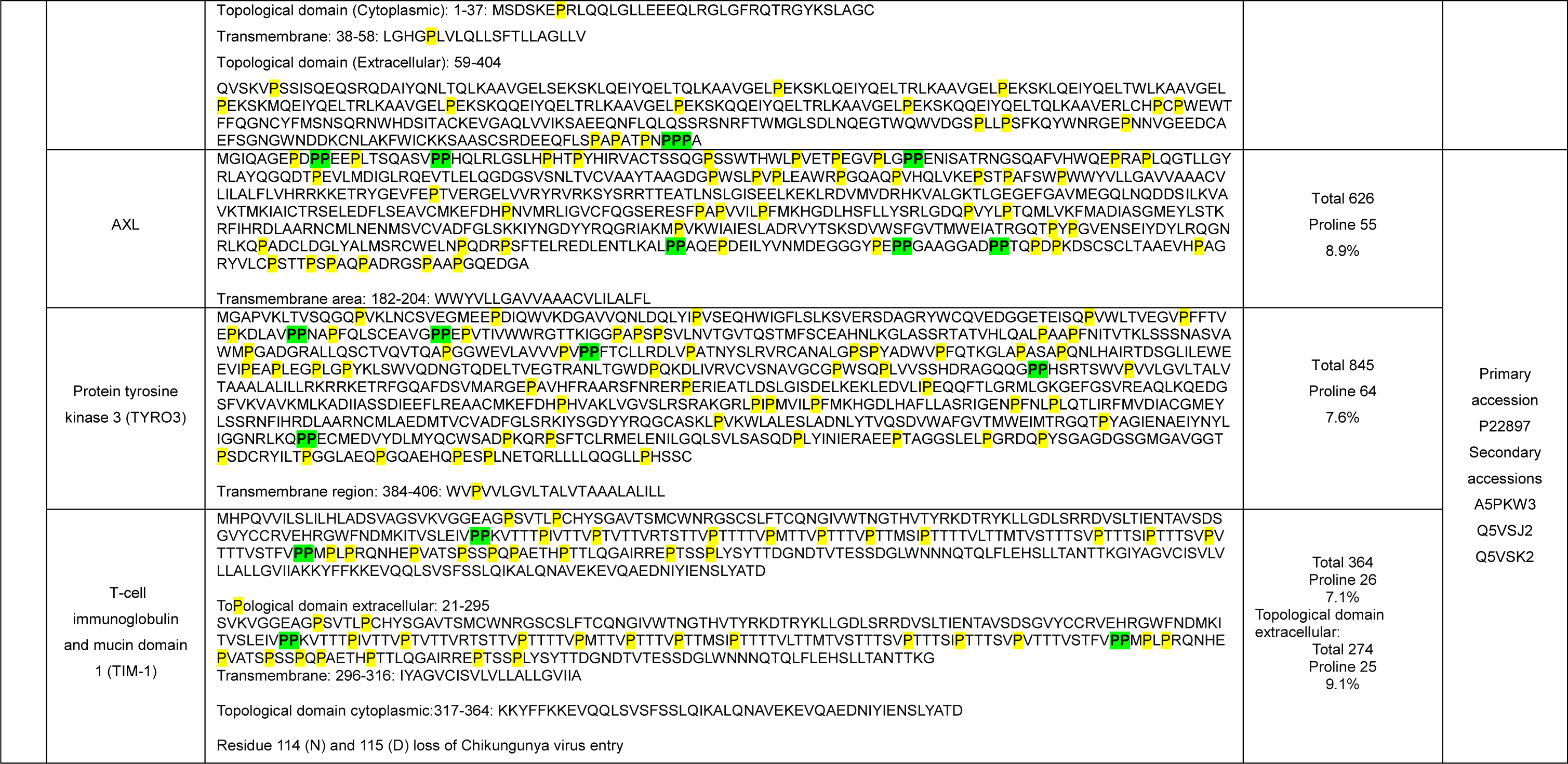
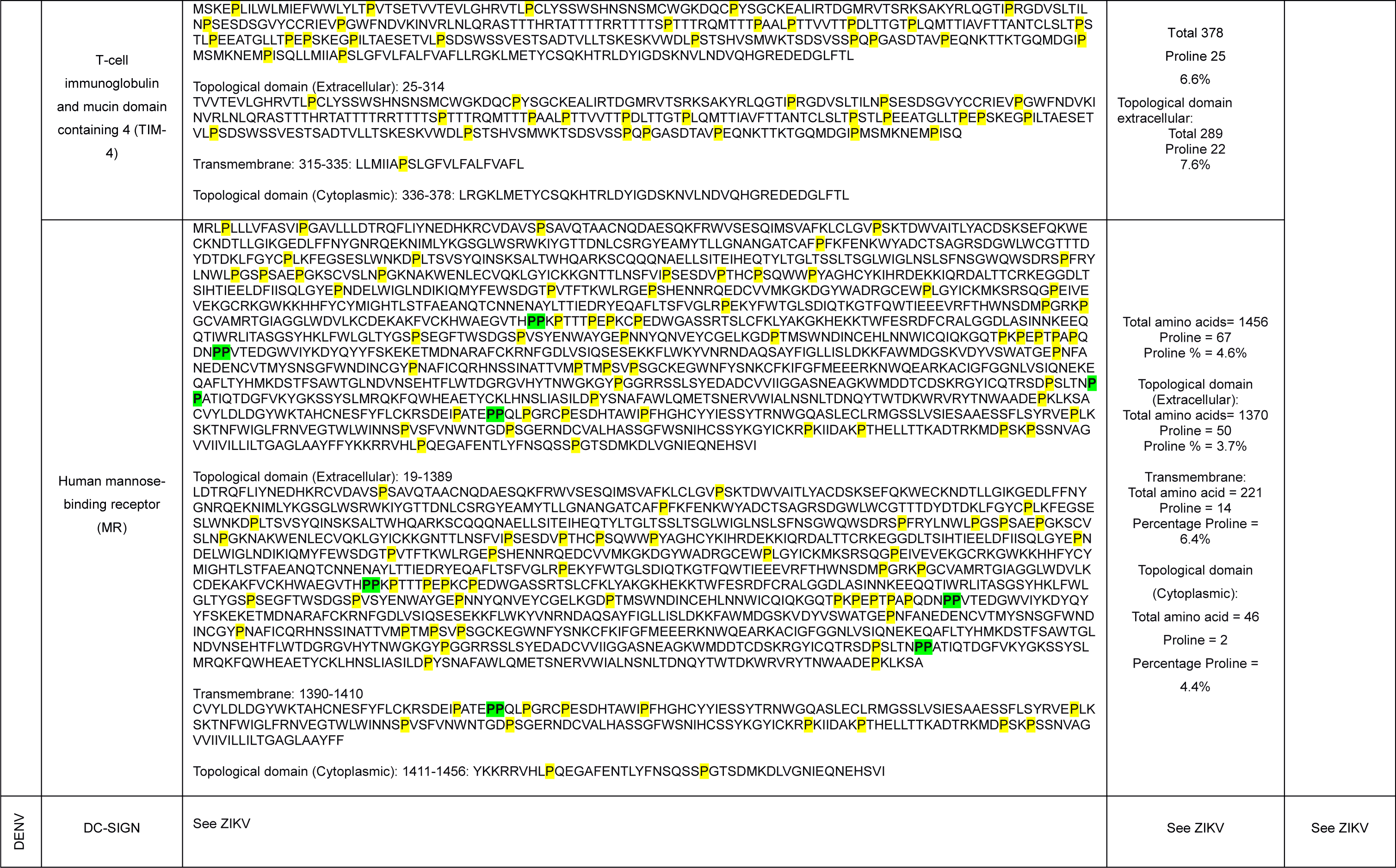
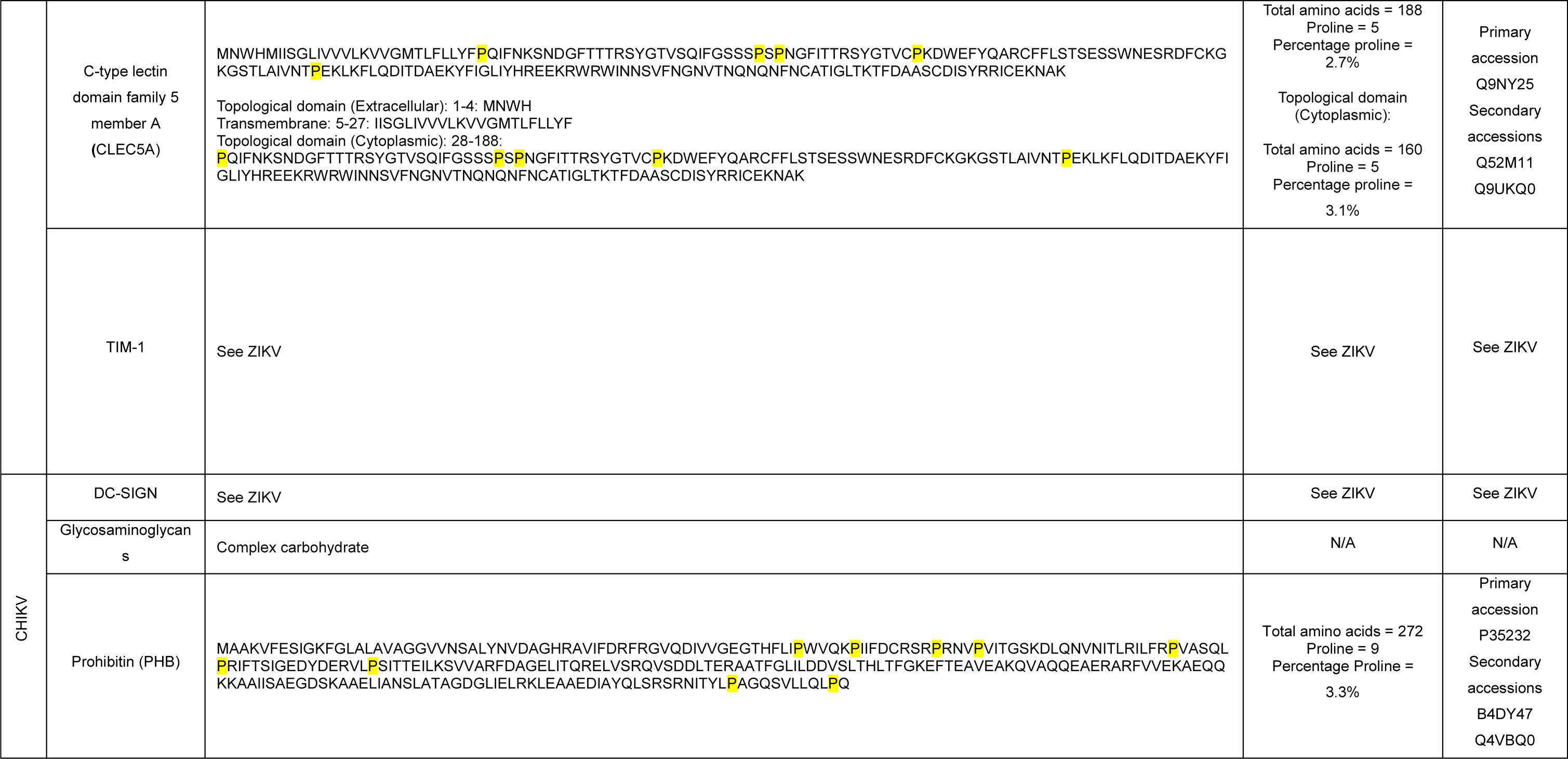
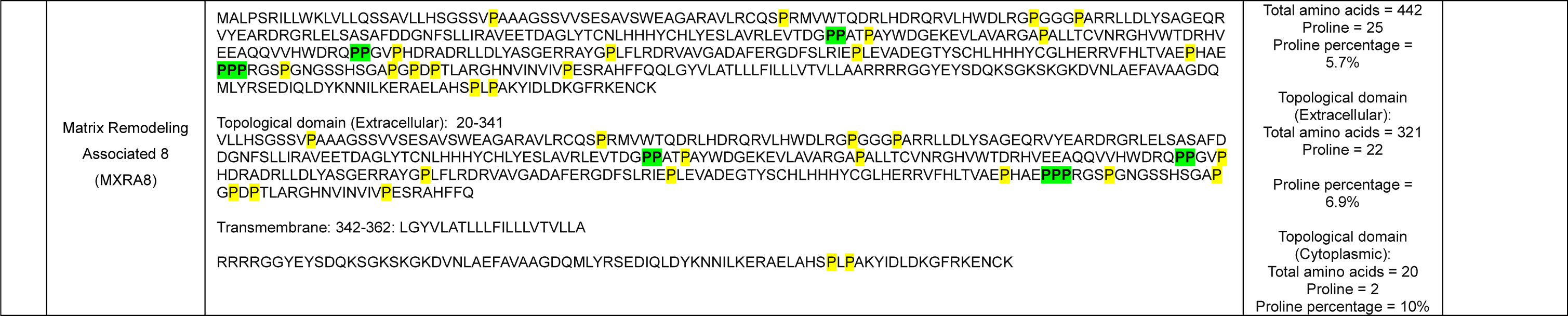
Estimated proline content and location of the important host factors for the viruses used in this study and viruses for which HH showed antiviral activity. Prolines are highlighted in yellow and polyproline motifs in green.

**Supplementary Table 5.**
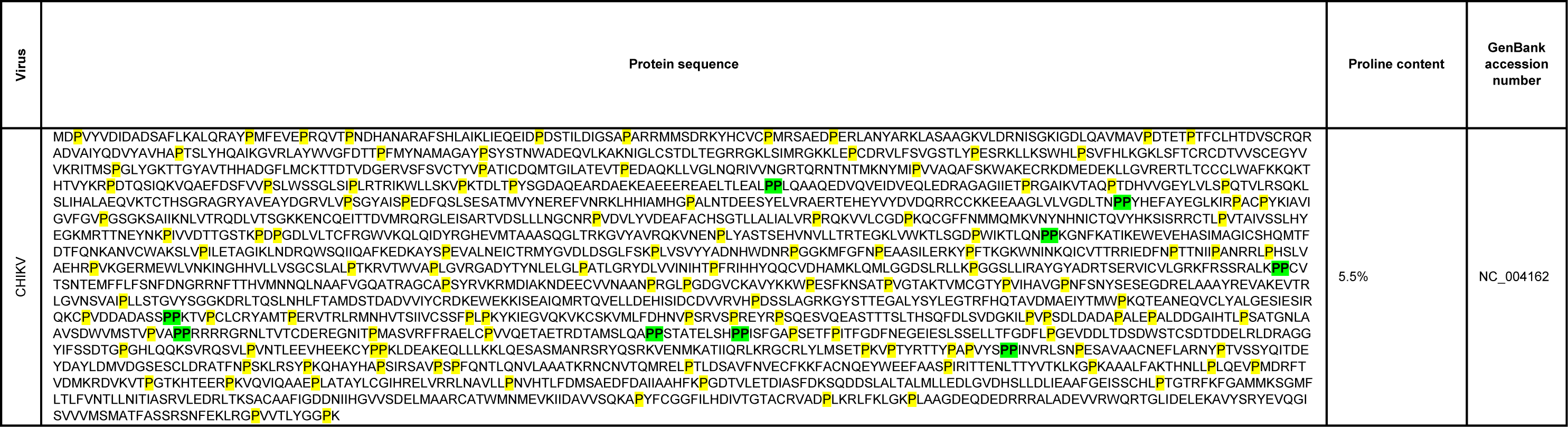
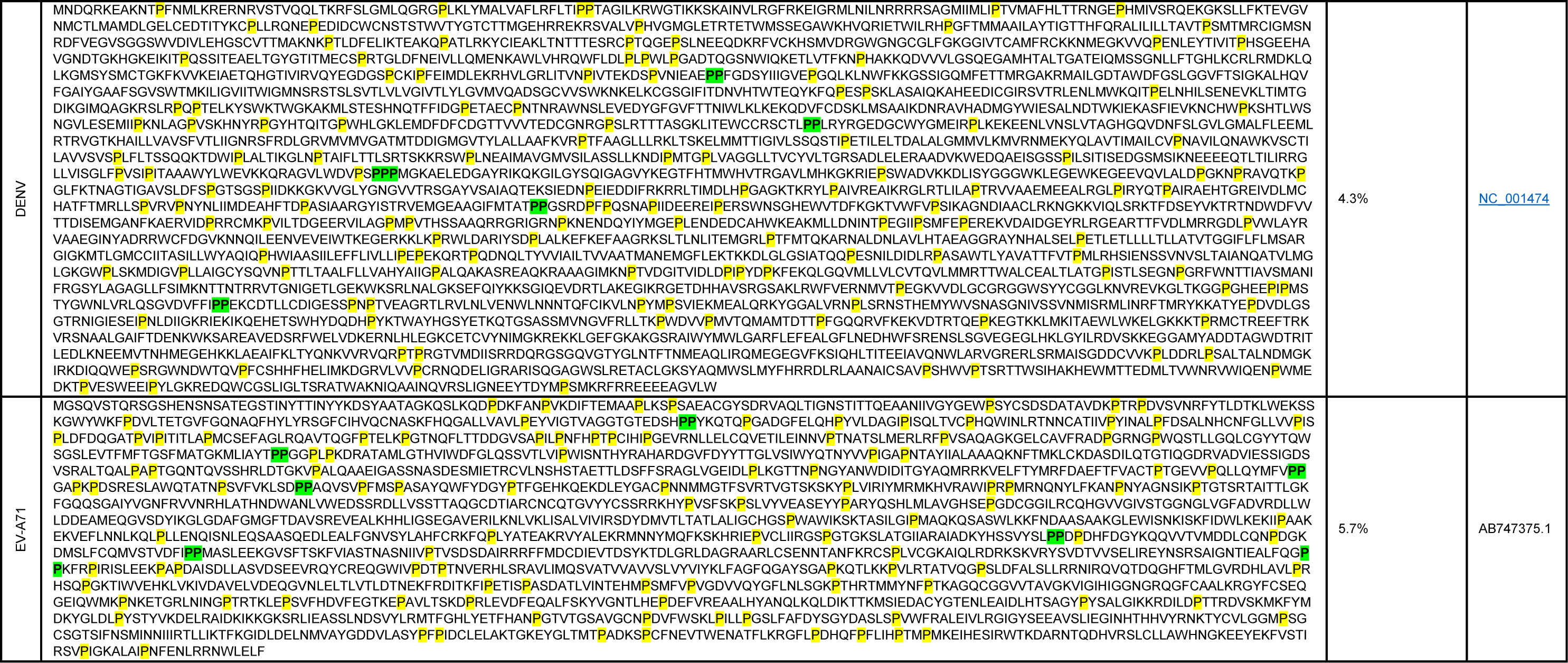
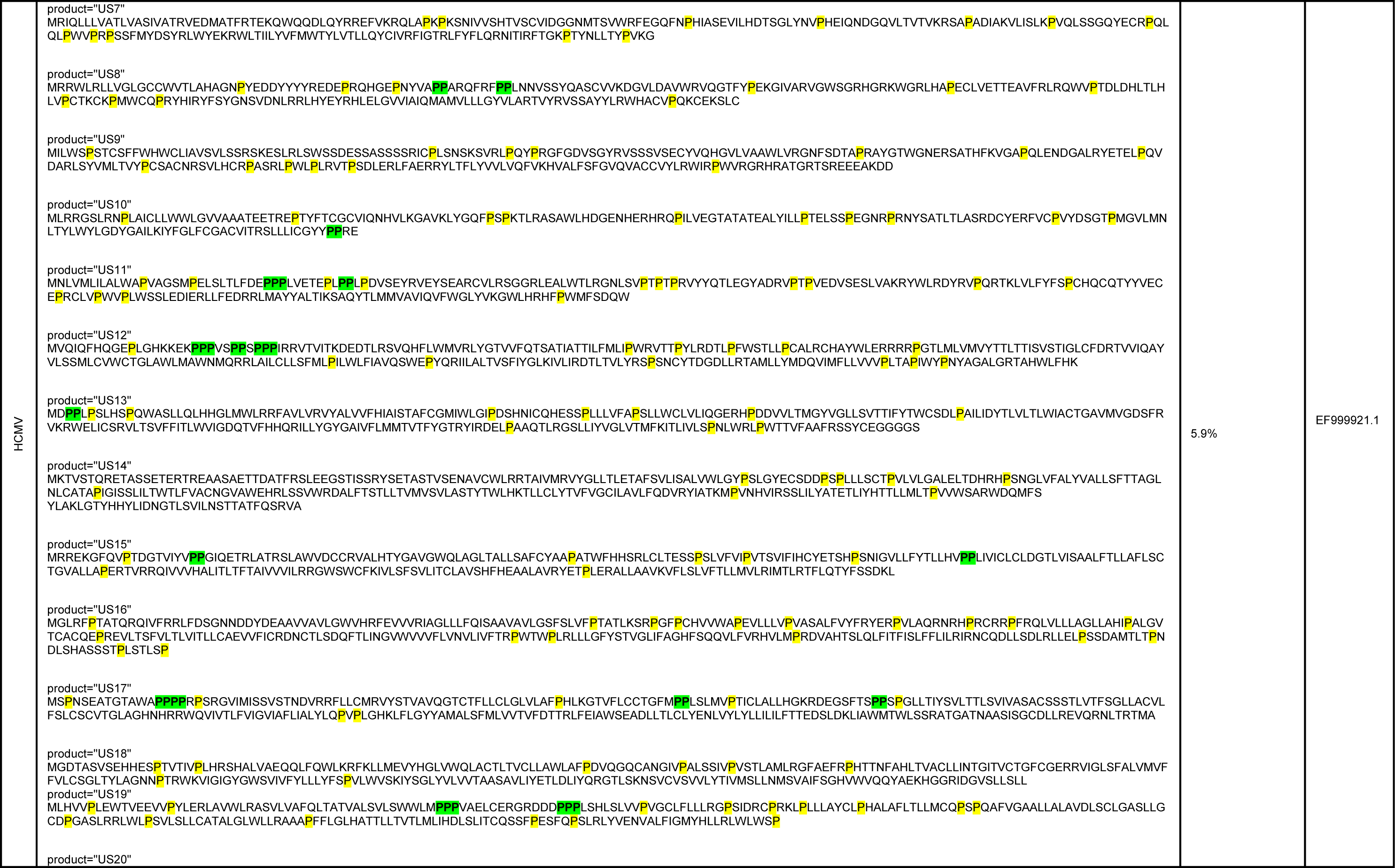
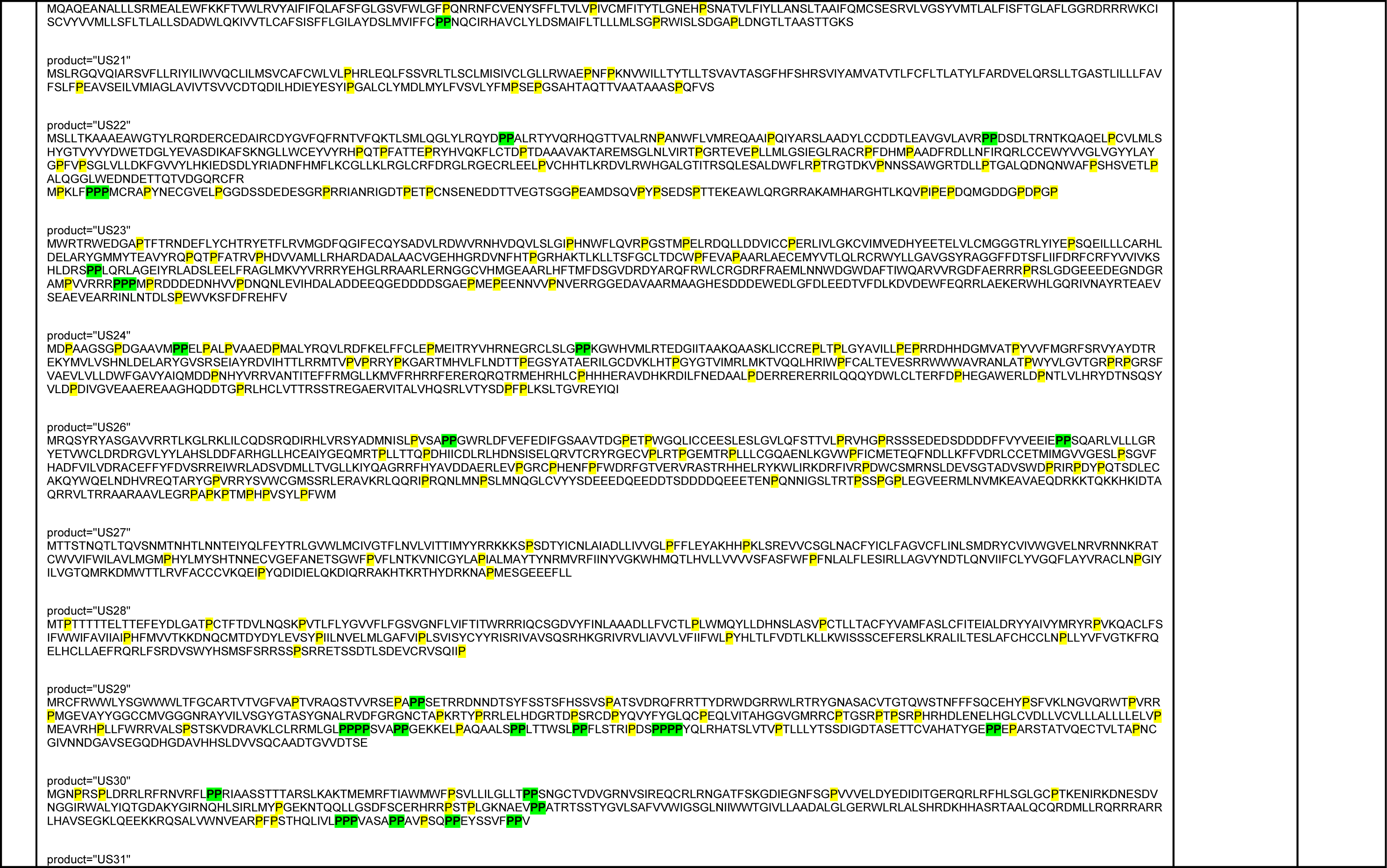
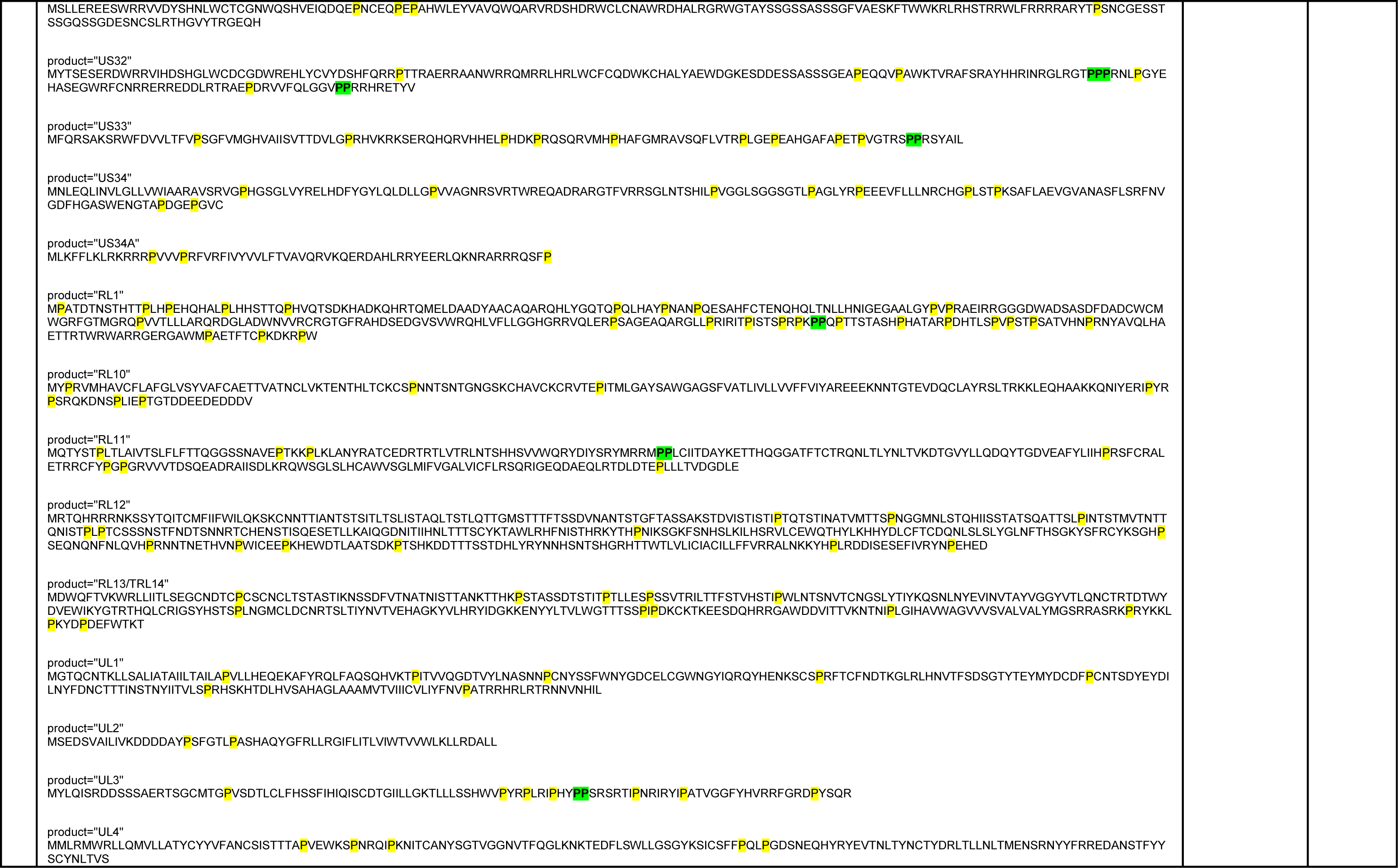
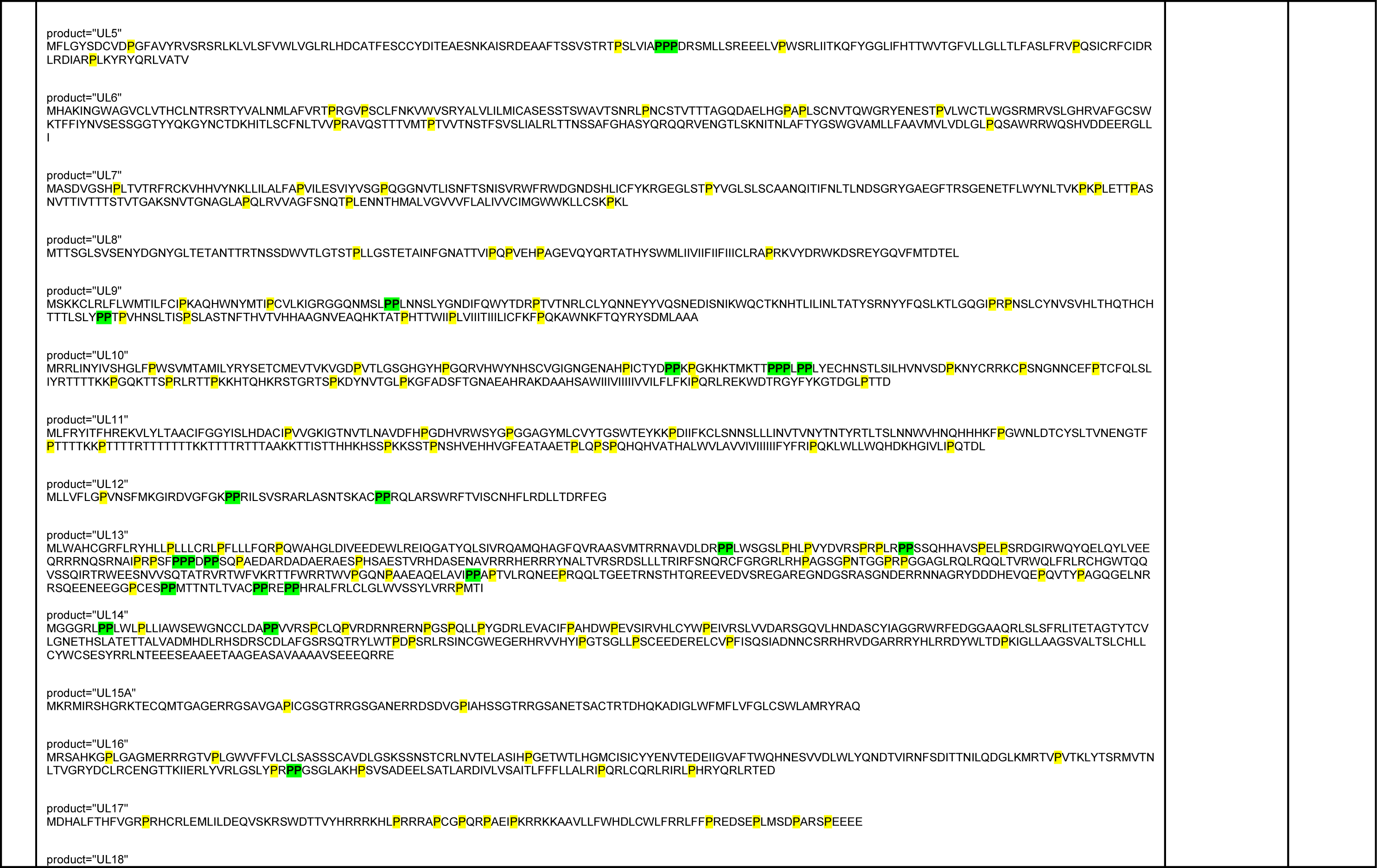
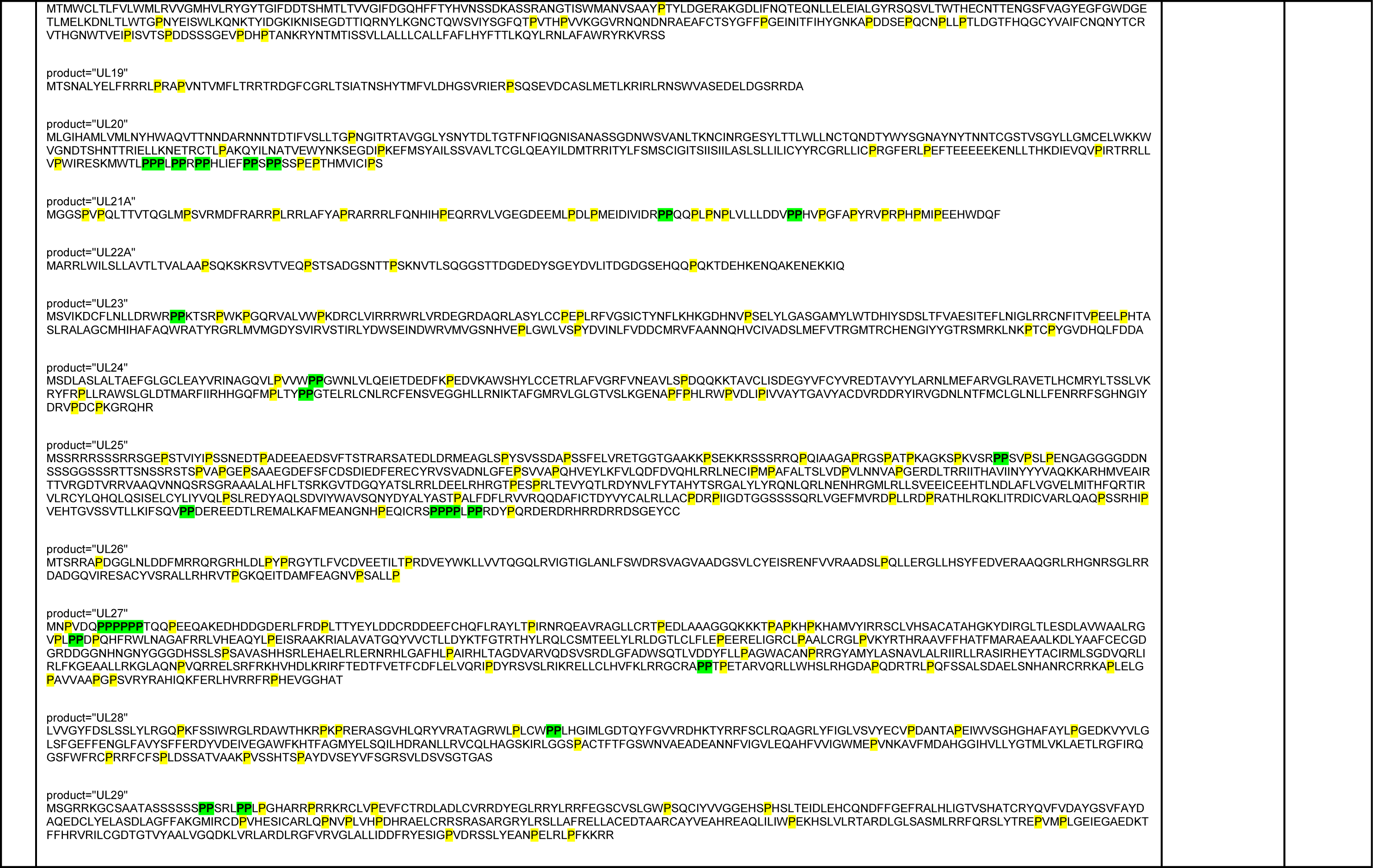
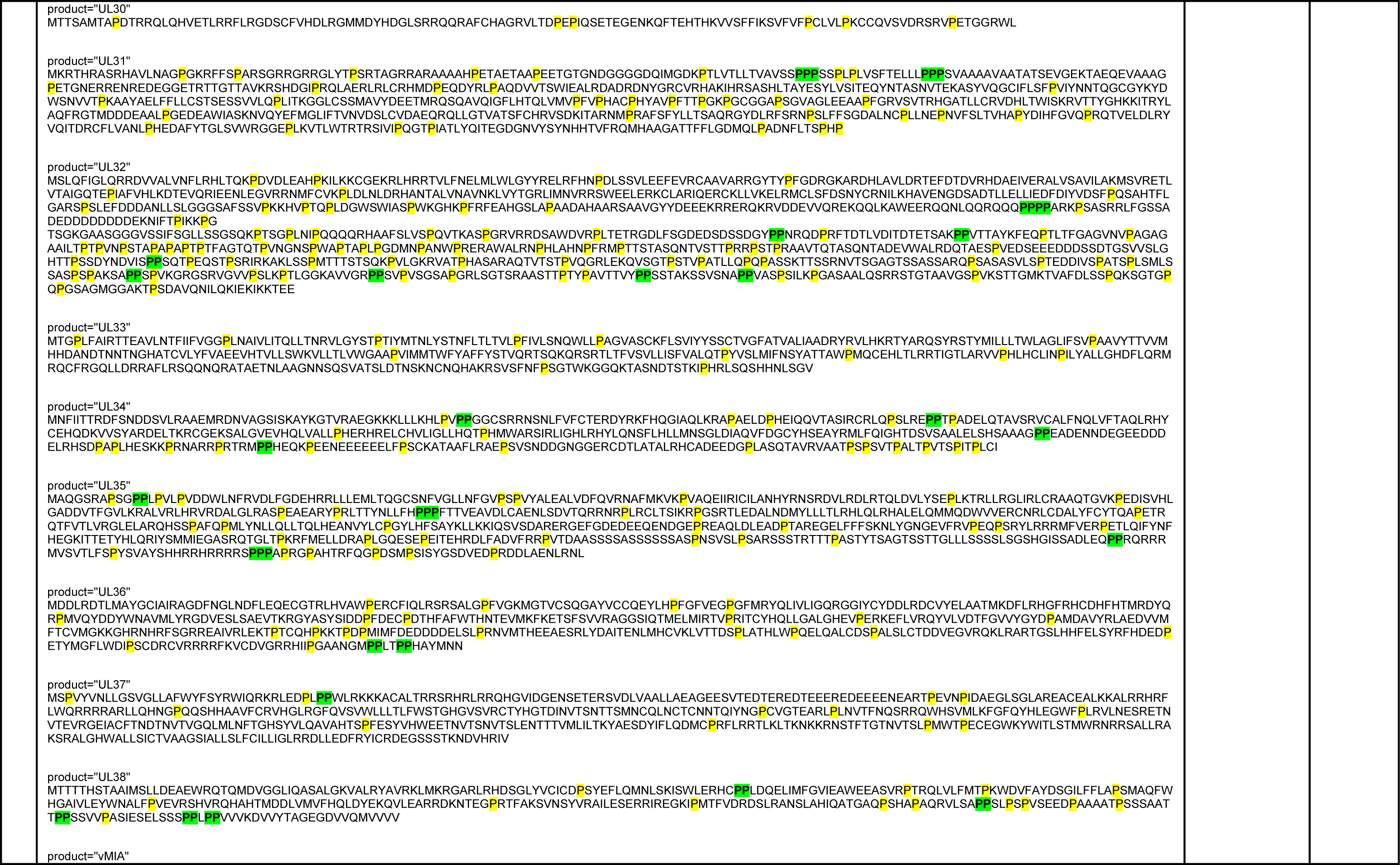
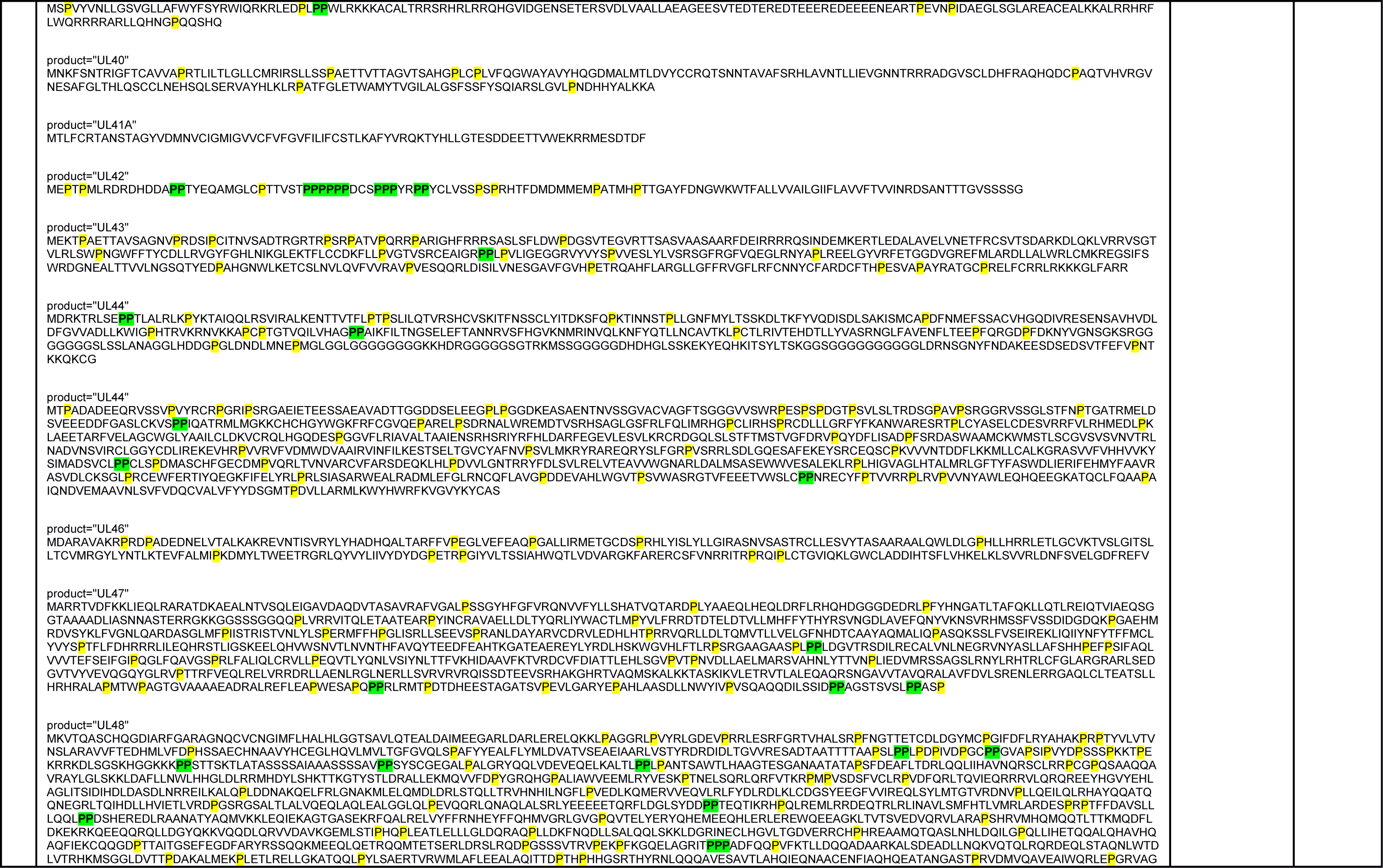
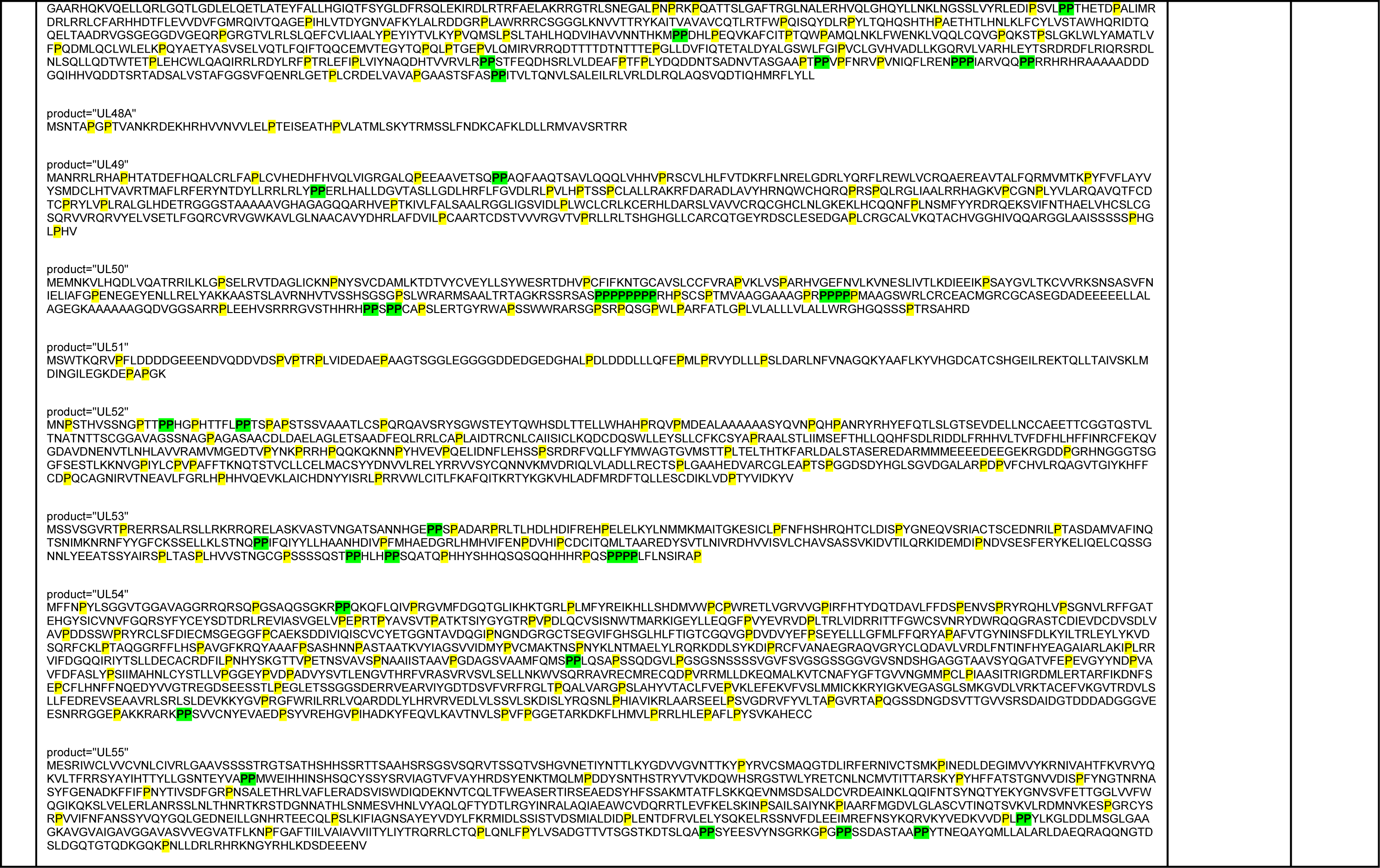
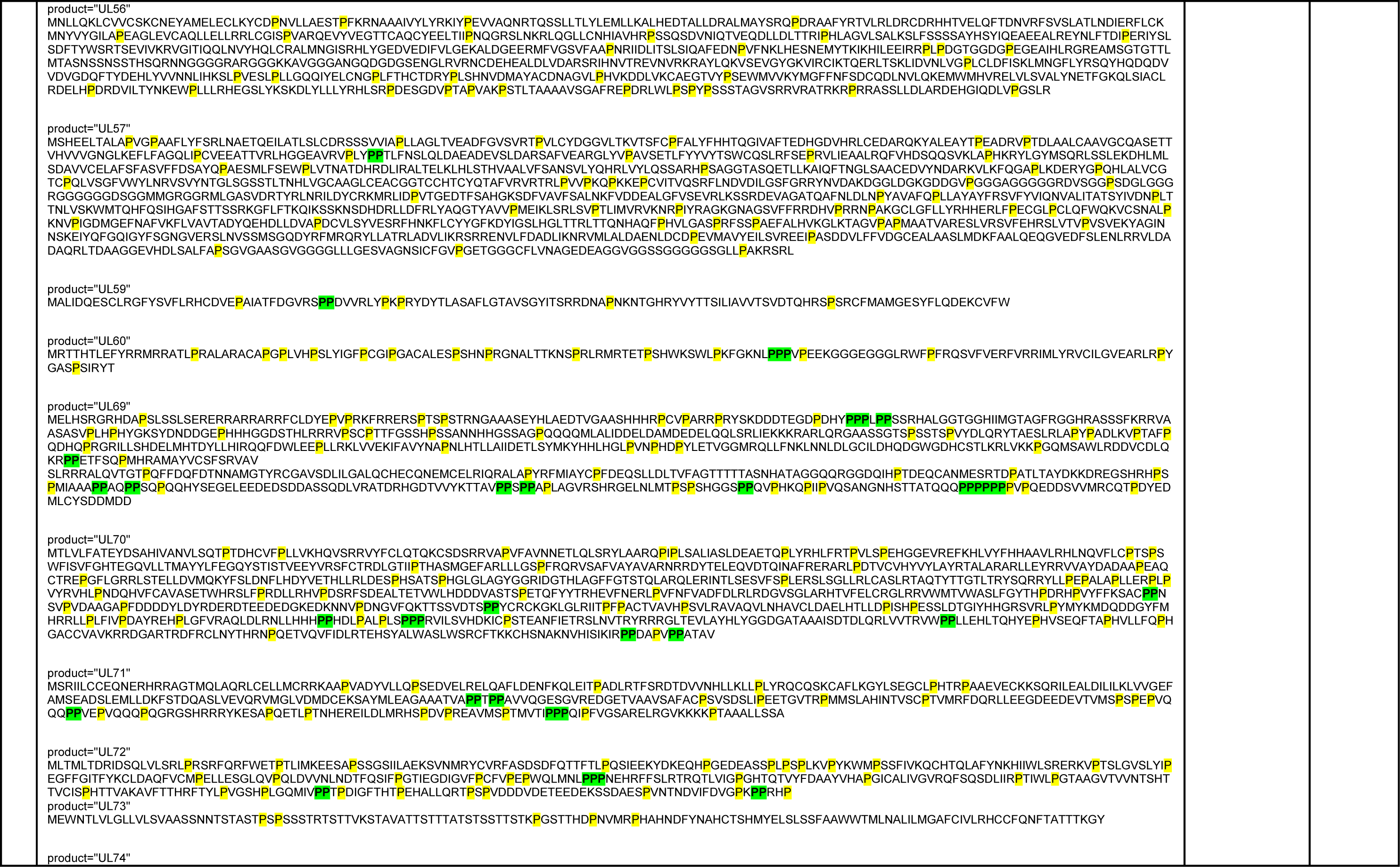
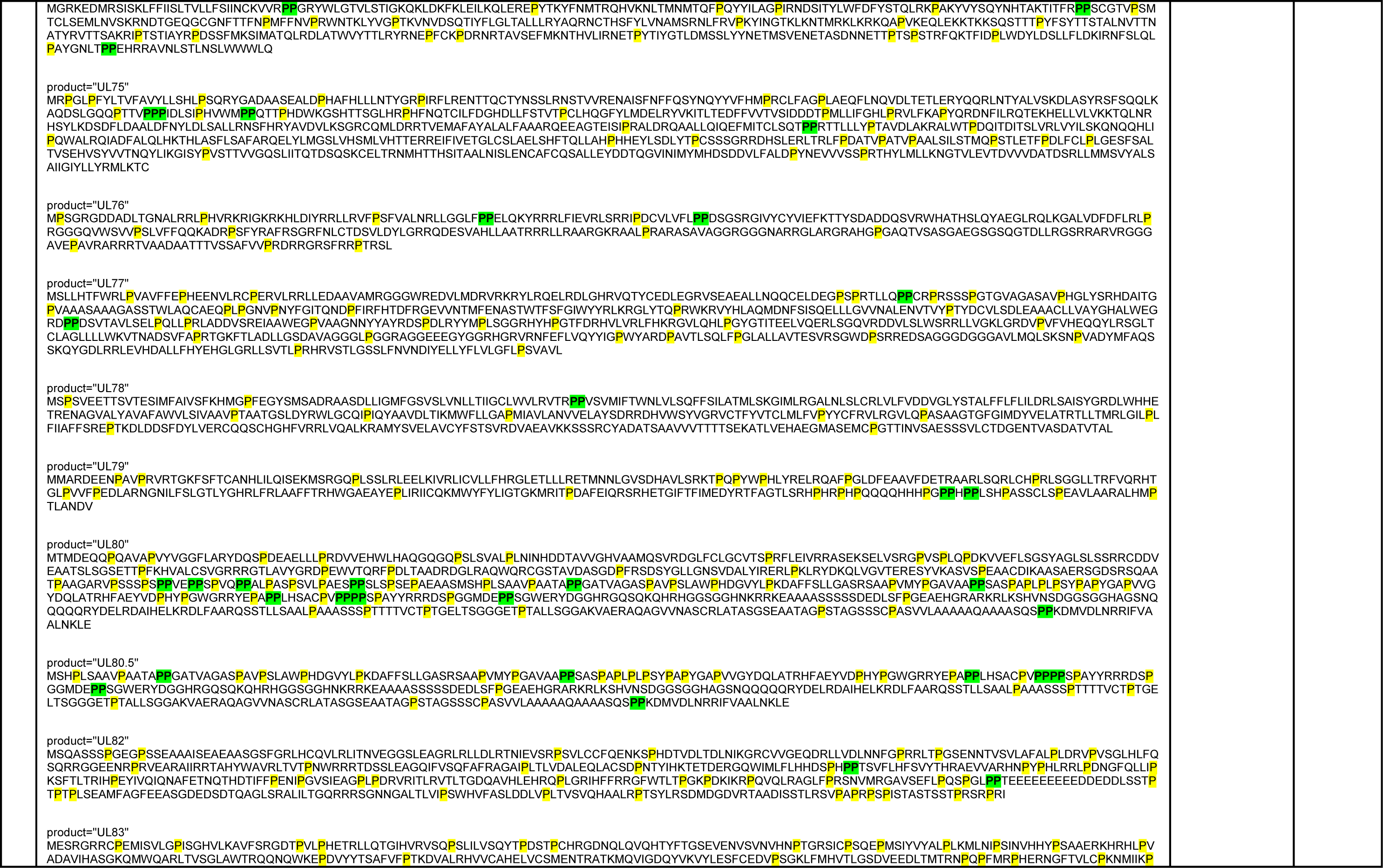
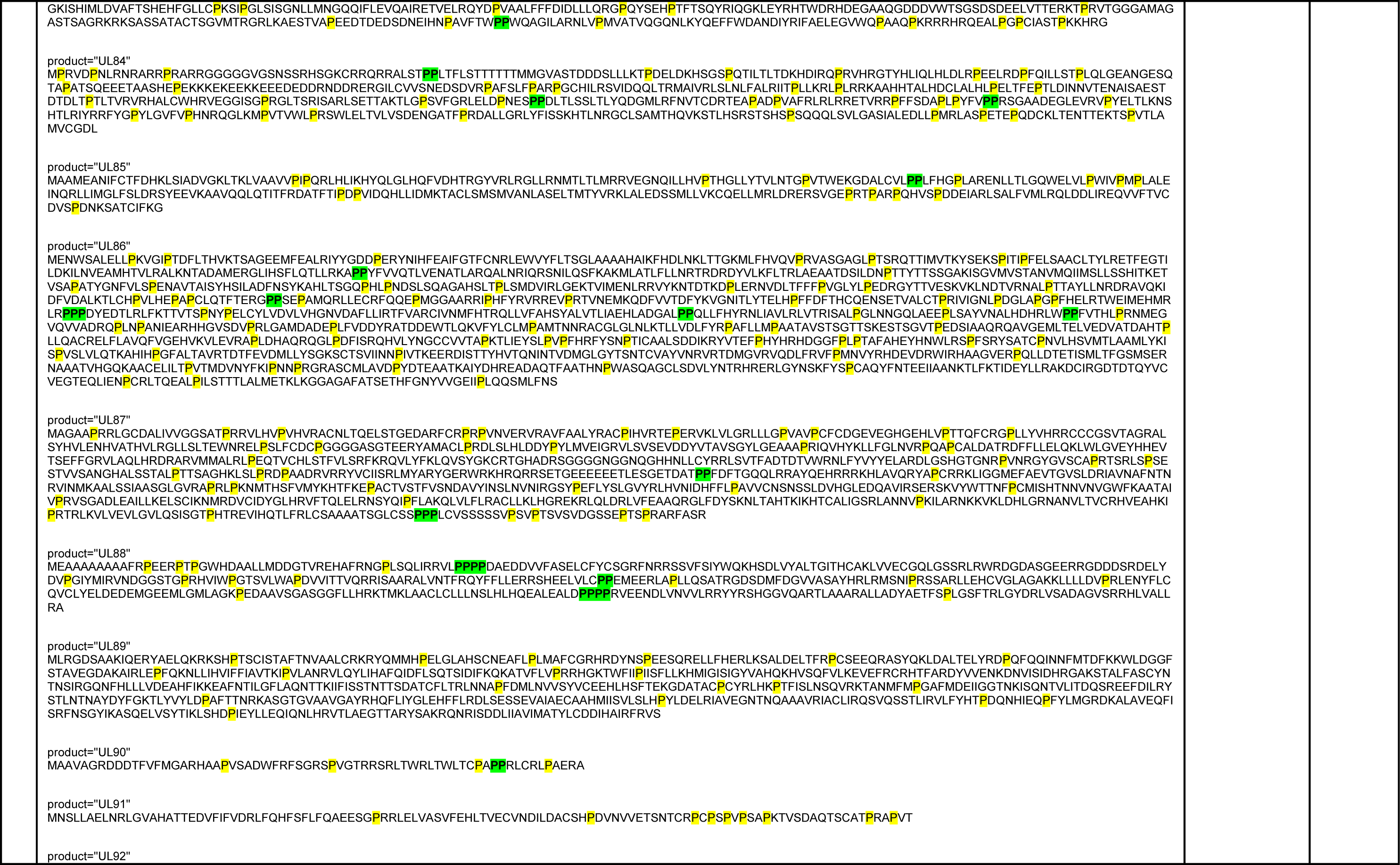
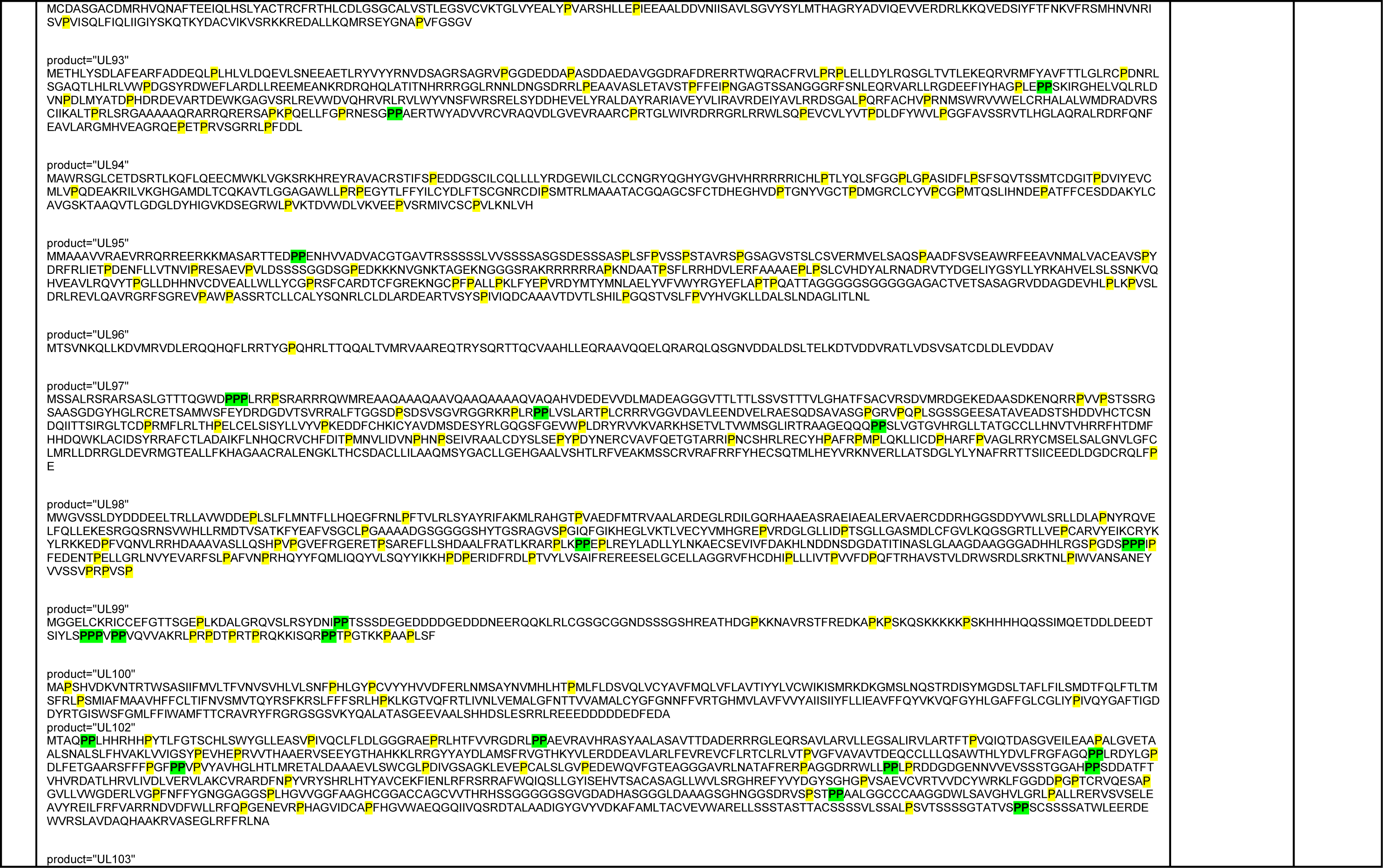
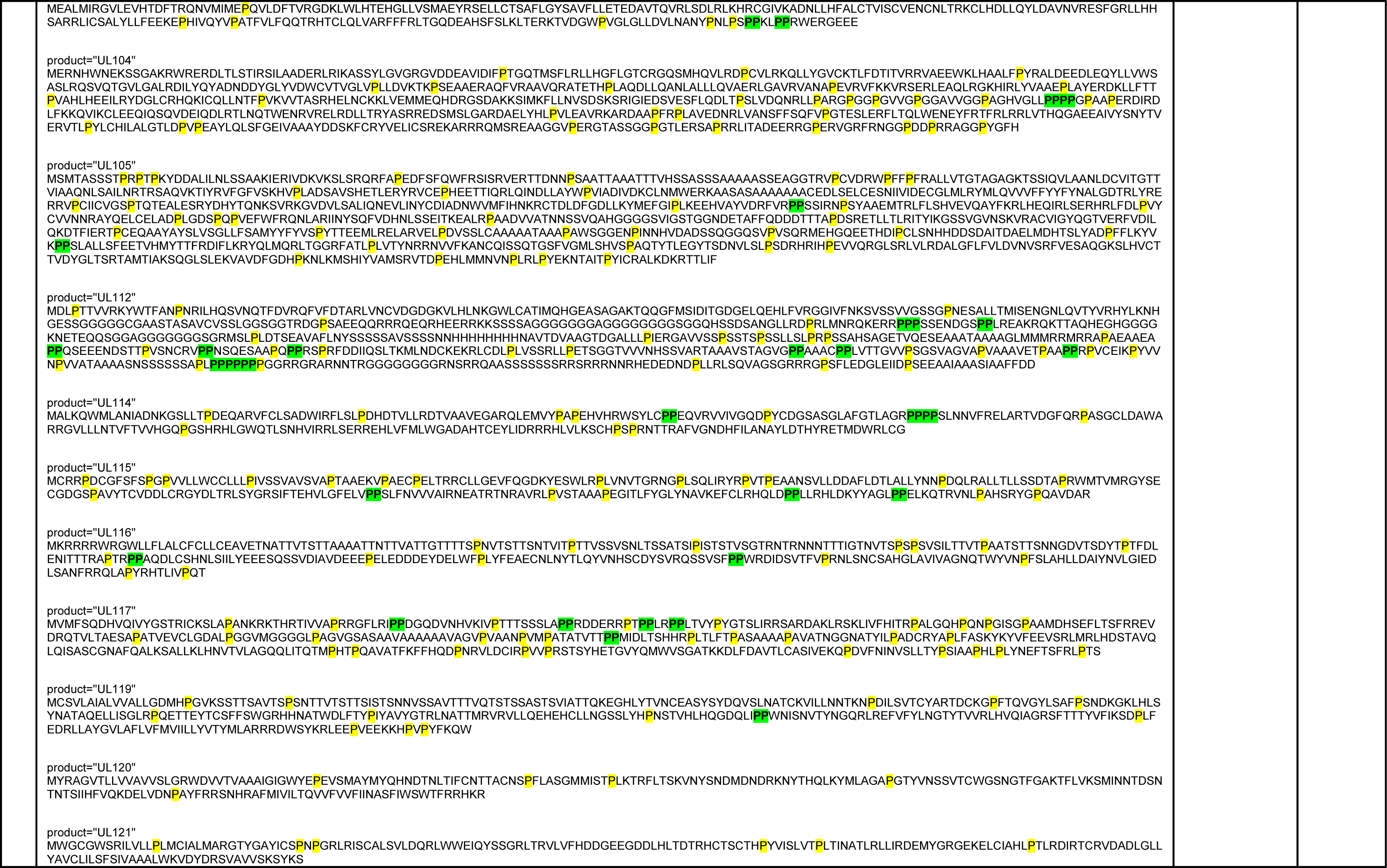
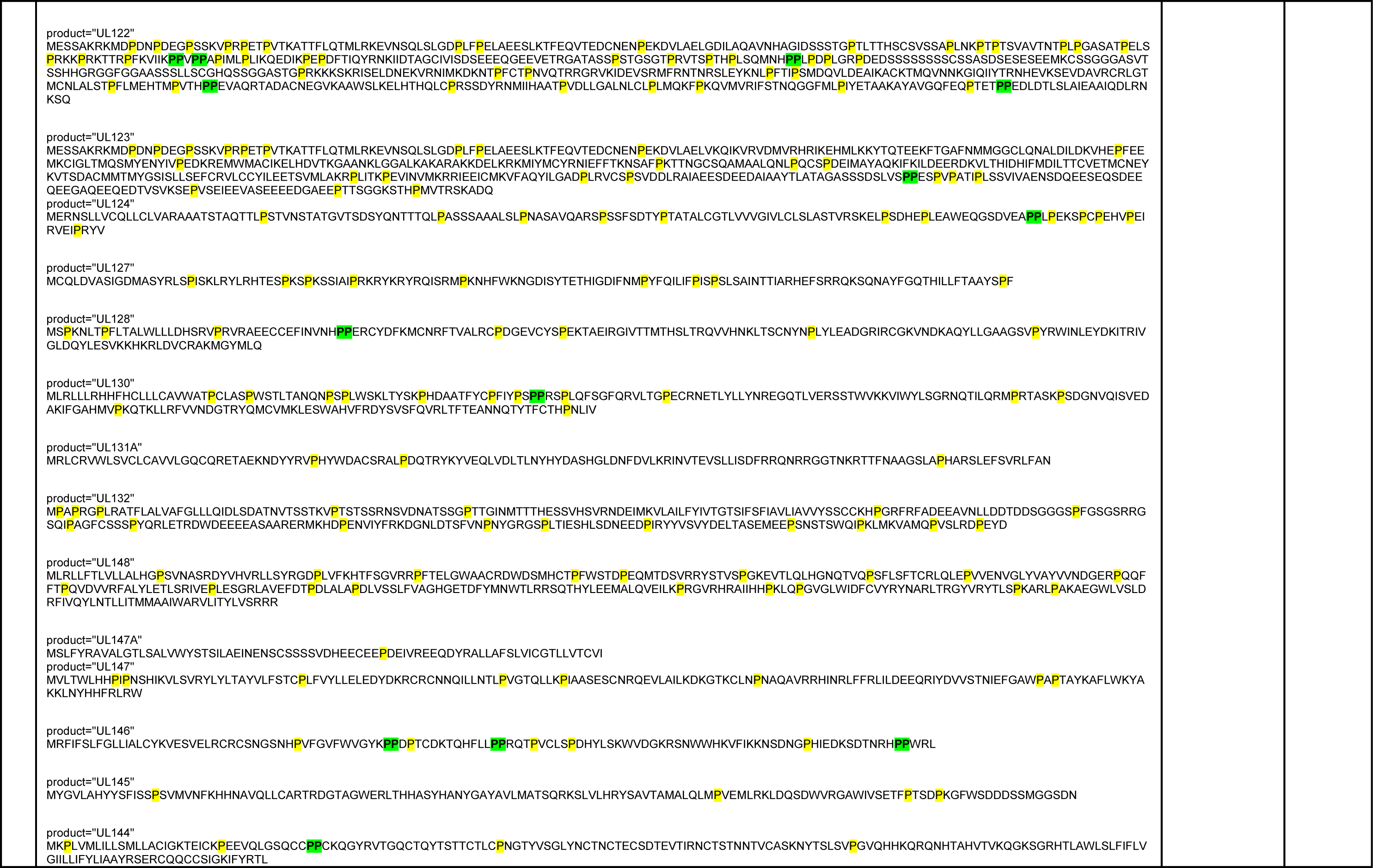
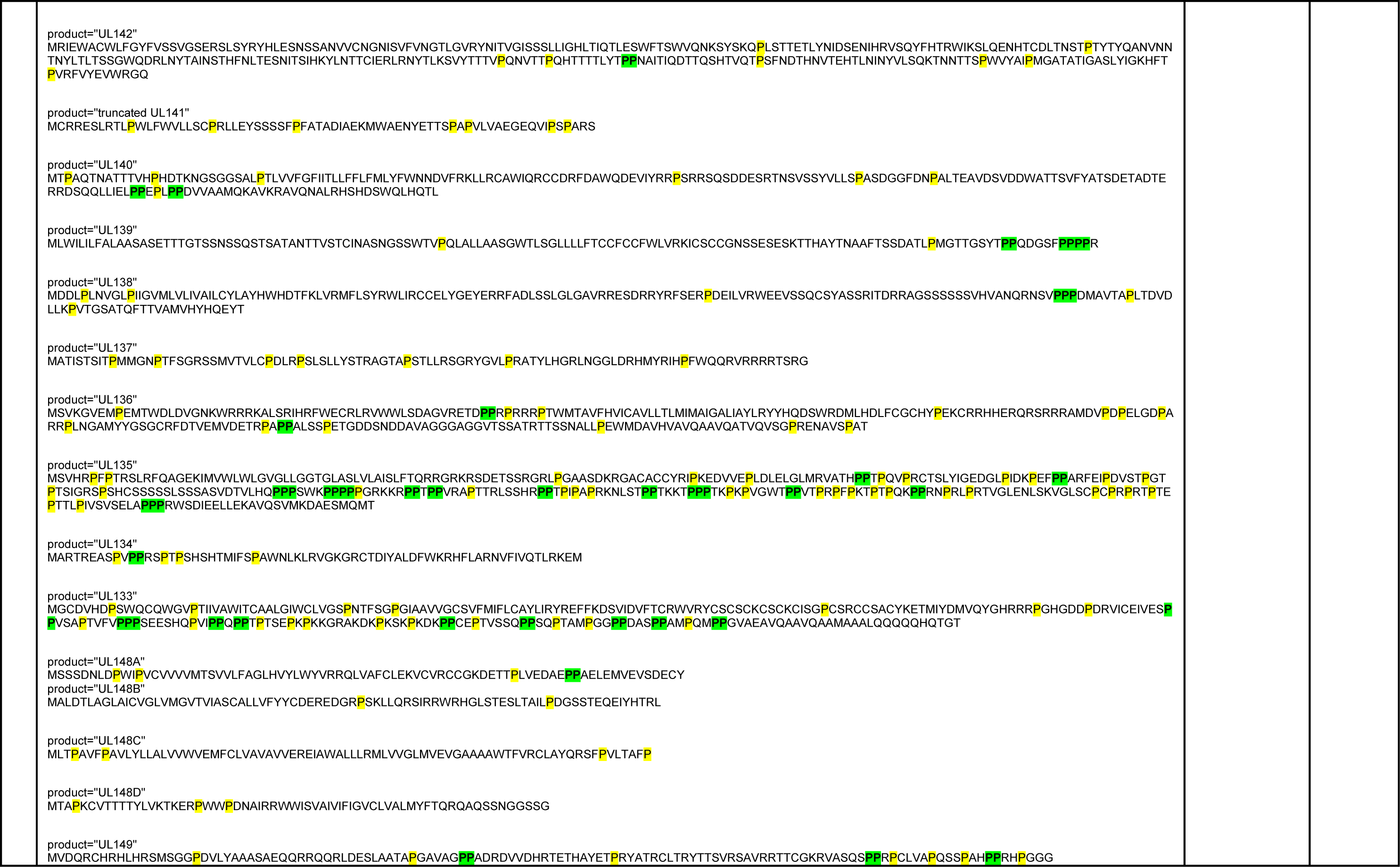
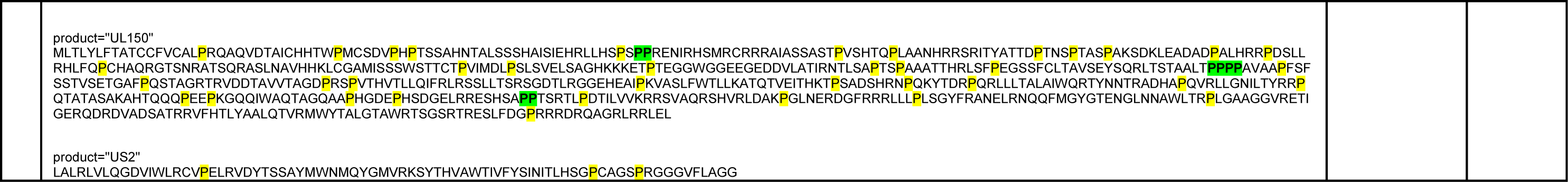
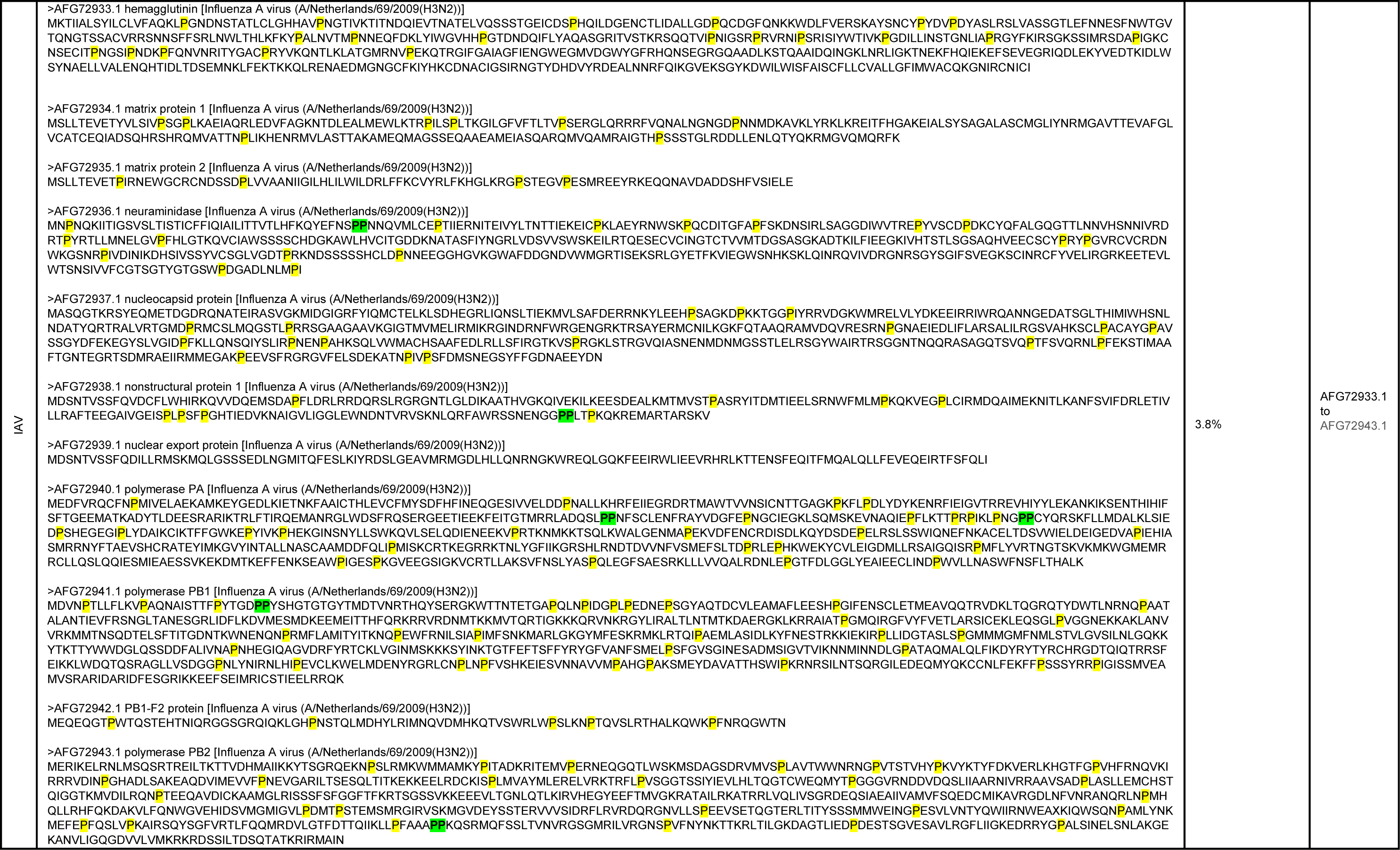
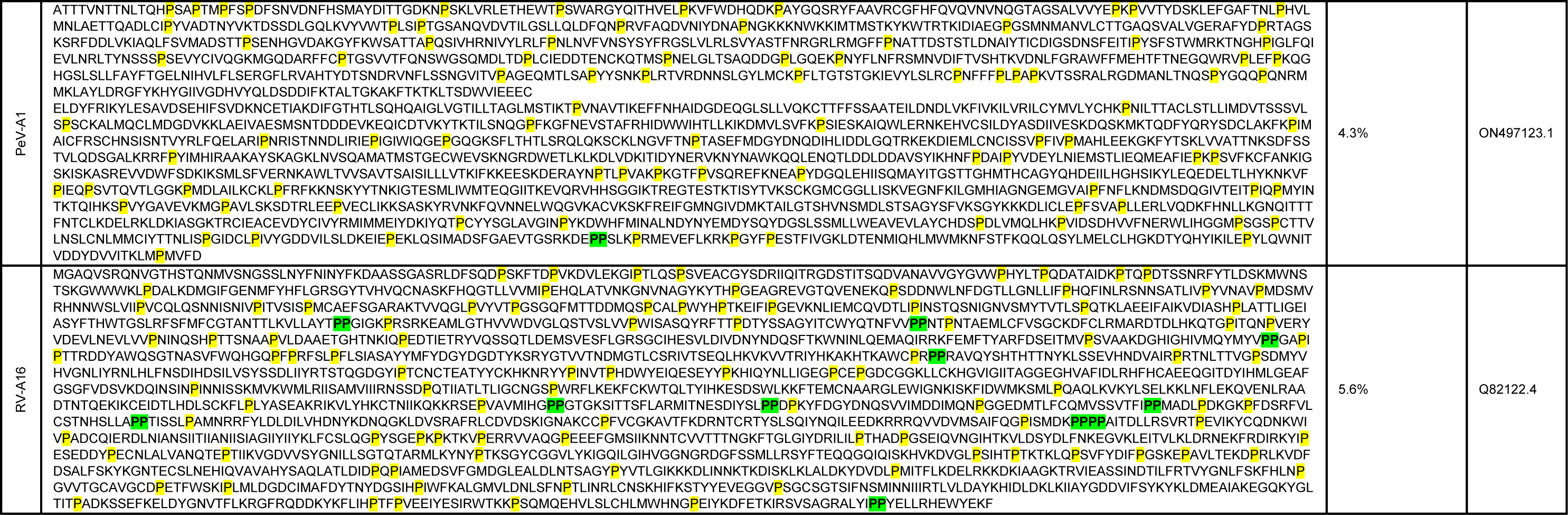
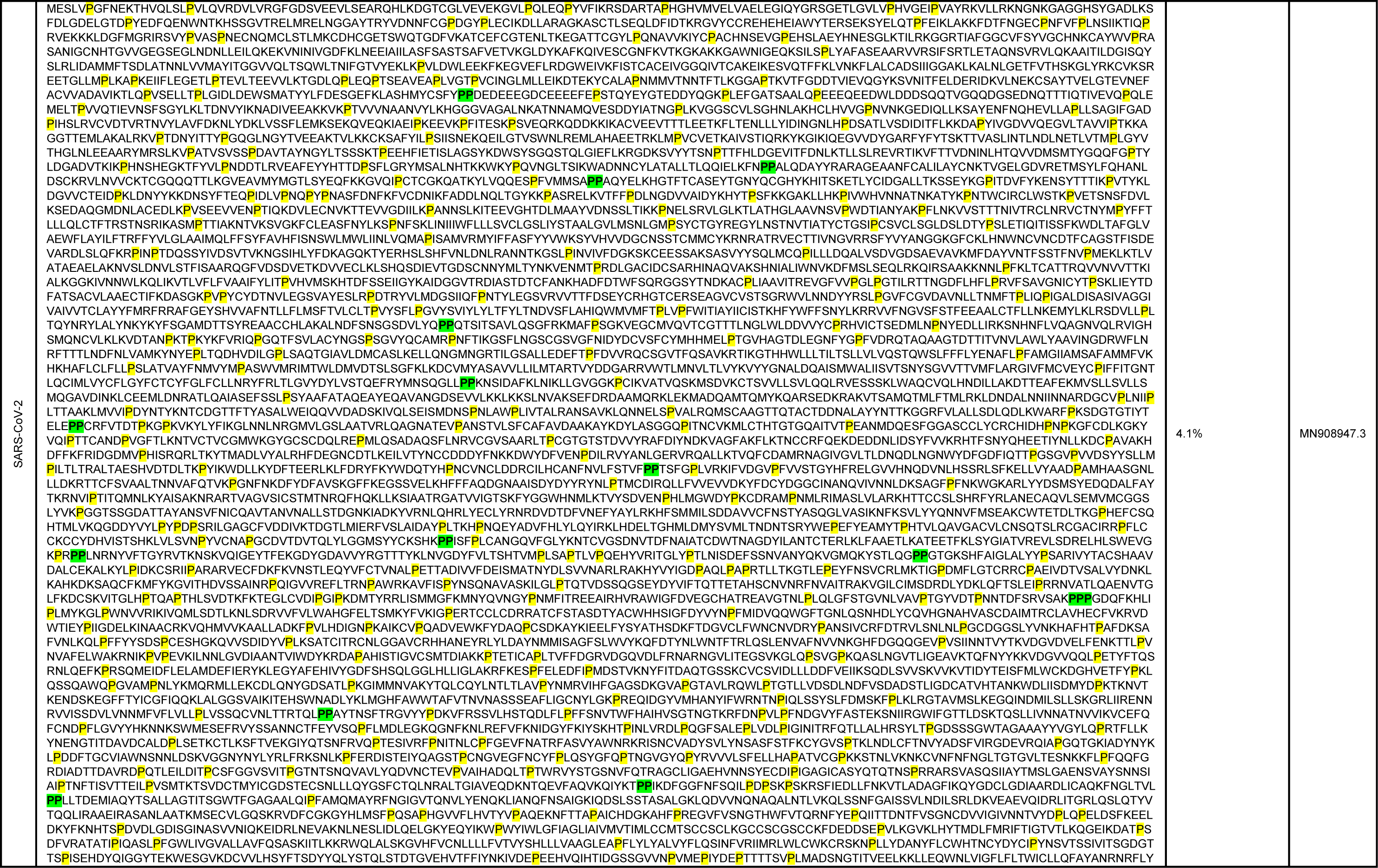
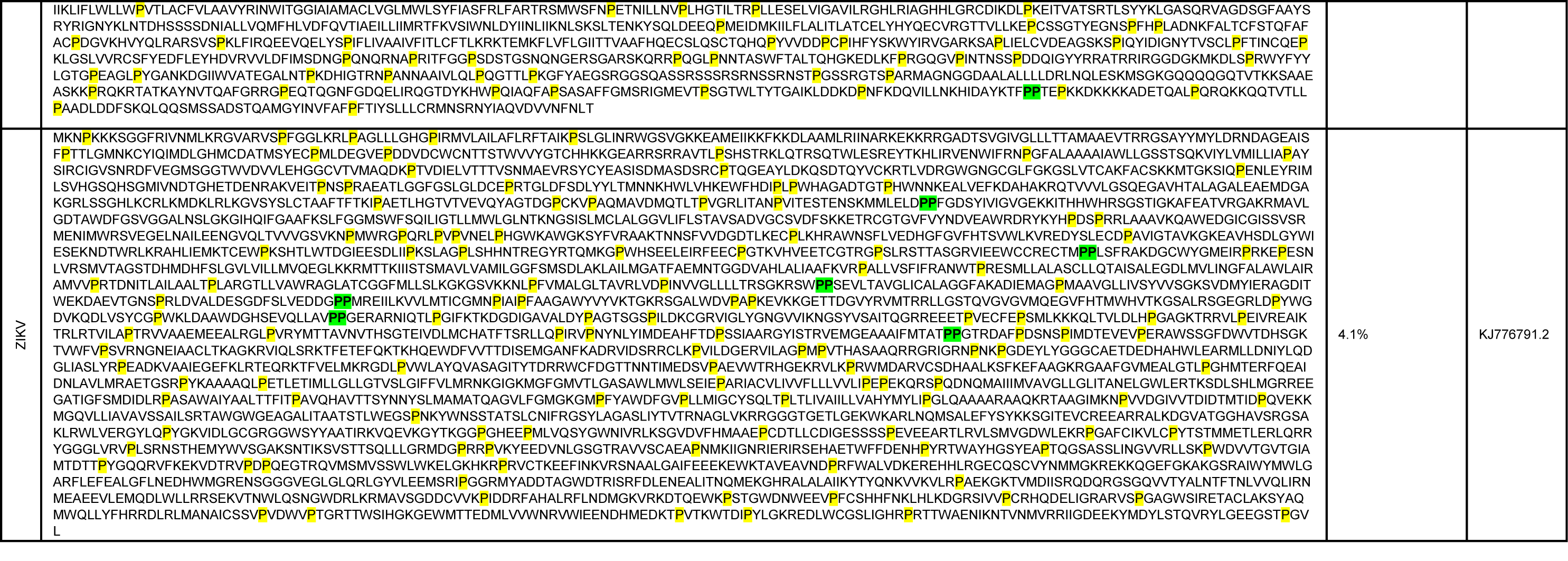
Estimated proline content of the viruses used in this study and viruses for which HH showed antiviral activity. Prolines are highlighted in yellow and polyproline motifs in green.

**Supplementary Figure 1.**
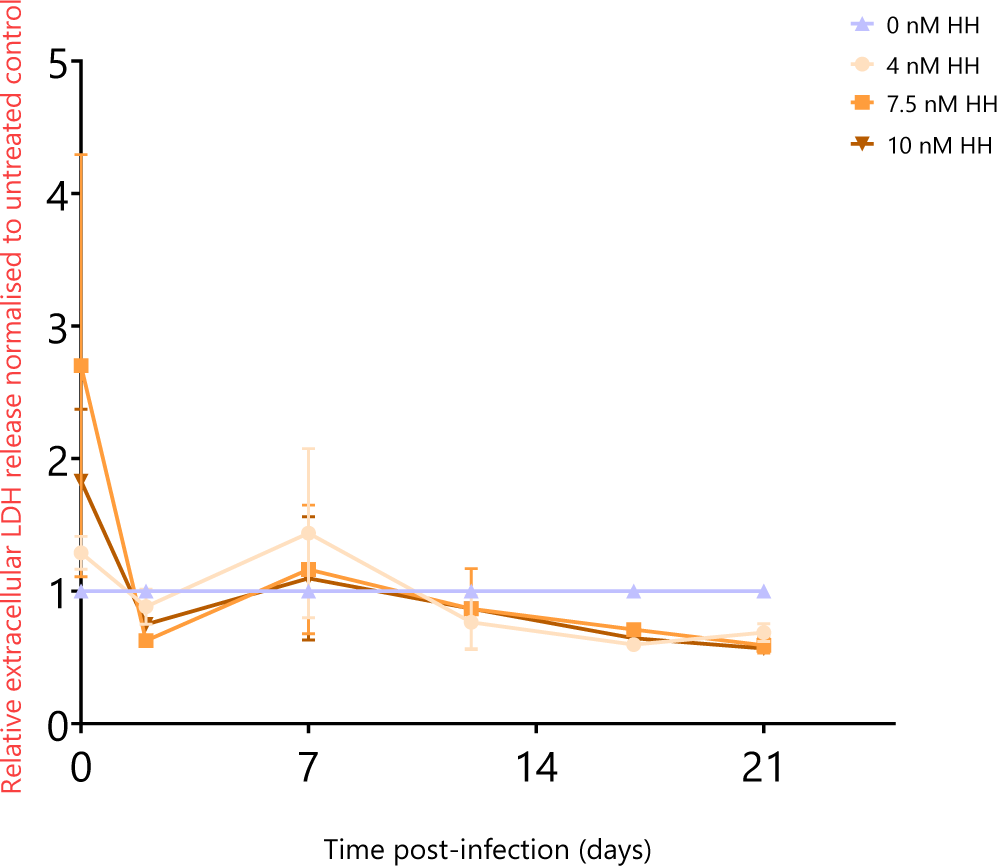
Relative LDH release normalized to untreated control in RNOs treated with different concentrations of HH refreshed on 2, 7, 12, and 17. Data represents the mean ± SEM two biological replicates in technical triplicates.

**Supplementary Figure 2.**
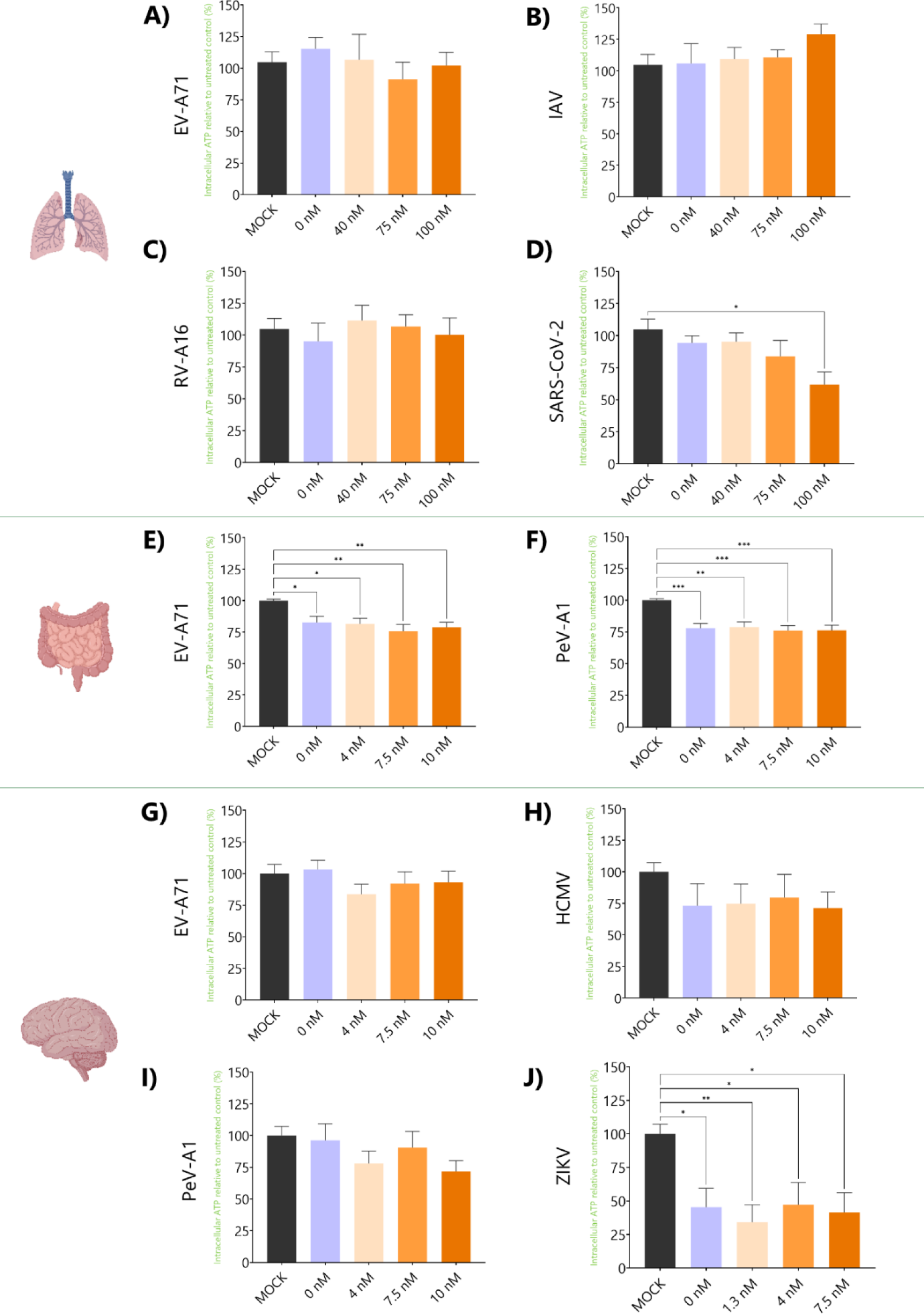
Intracellular ATP relative to untreated controls (%) in HAE infected with EV-A71 (A), IAV (B), RV-A16 (C), SARS-CoV-2 (E); or HIE infected with EV-A71 (E), PeV-A1 (F); or RNOs infected with EV-A71 (G), HCMV (H), PeV-A1 (I), ZIKV (J) and treated with different concentrations of HH. In all cases, data represents the mean ± SEM of HAE/ HIE three biological replicates in technical duplicates, and RNOs three biological replicates in technical triplicates. Statistical significance was determined using One-way ANOVA with Tukey’s multiple comparisons. * p-value < 0.05; ** p-value <0.01; *** p-value <0.001.

